# Empirical Transmit Field Bias Correction of T1w/T2w Myelin Maps

**DOI:** 10.1101/2021.08.08.455570

**Authors:** Matthew F. Glasser, Timothy S. Coalson, Michael P. Harms, Junqian Xu, Graham L. Baum, Joonas A. Autio, Edward J. Auerbach, Douglas N. Greve, Essa Yacoub, David C. Van Essen, Nicholas A. Bock, Takuya Hayashi

## Abstract

T1-weighted divided by T2-weighted (T1w/T2w) myelin maps were initially developed for neuroanatomical analyses such as identifying cortical areas, but they are increasingly used in statistical comparisons across individuals and groups with other variables of interest. Existing T1w/T2w myelin maps contain radiofrequency transmit field (B1+) biases, which may be correlated with these variables of interest, leading to potentially spurious results. Here we propose two empirical methods for correcting these transmit field biases using either explicit measures of the transmit field or alternatively a ‘pseudo-transmit’ approach that is highly correlated with the transmit field at 3T. We find that the resulting corrected T1w/T2w myelin maps are both better neuroanatomical measures (e.g., for use in cross-species comparisons), and more appropriate for statistical comparisons of relative T1w/T2w differences across individuals and groups (e.g., sex, age, or body-mass-index) within a consistently acquired study at 3T. We recommend that investigators who use the T1w/T2w approach for mapping cortical myelin use these B1+ transmit field corrected myelin maps going forward.

## 1. Introduction

The T1-weighted divided by T2-weighted (T1w/T2w) myelin mapping approach was originally developed over a decade ago as a high-resolution non-invasive measure of cortical architecture that correlates with postmortem measures of cortical myelin content for the purpose of mapping cortical areas (though the relationship of the T1w/T2w ratio to myelin content of subcortical structures in the volume is more complex; Glasser and Van Essen 2011). The T1w/T2w approach has proven highly successful at mapping cortical areas, revealing many cortical areal boundaries that co-localize with those of other non-invasive measures of cortical properties, such as cortical thickness, resting state functional connectivity, task activation, and within-area topographic maps measured with functional connectivity (Glasser et al., 2016a). Initial efforts were focused at the group level, given artifacts from imperfect surface reconstructions in individuals from relatively low resolution 1mm isotropic T1w and T2w scans (Glasser and Van Essen 2011). Markedly improved individual participant T1w/T2w myelin maps (Glasser et al., 2013; 2014) were later achieved by improving spatial resolution to 0.7mm or 0.8mm isotropic, less than half of the minimum 1.6mm cortical thickness (Glasser et al., 2016b), and improving FreeSurfer surface reconstructions (Fischl 2012; Zaretskaya et al., 2018) by using the full resolution of the T1w images for white and pial surface placement and using the T2w image for exclusion of dura and blood vessels during pial surface placement. These individual myelin maps, collected and processed by the Human Connectome Project (HCP), were of sufficient quality to enable individual participant identification of cortical areas using a machine learning algorithm when combined with the other multi-modal non-invasive measures mentioned above (Glasser et al., 2016a). Importantly, the methodological improvements in T1w/T2w myelin maps in the context of the HCP Pipelines have to date focused solely on ensuring that they are appropriate for the above neuroanatomical use cases.

In parallel, there has been increasing interest in using non-invasive myelin maps, including T1w/T2w myelin maps, to make comparisons across individuals and groups, for example in healthy adults (Teraguchi et al., 2014; Shafee et al., 2015; Yang et al., 2020), over the course of development (Bozek et al., 2018; Grydeland et al., 2019; Norbom et al., 2020; Kwon et al., 2020), aging (Grydeland et al., 2013; 2019; Vidal-Piñeiro et al, 2016), in brain diseases (Teraguchi et al., 2014; Granberg et al., 2017; Rowley et al 2018; Nakamura et al., 2017; Du et al., 2019; Wei et al., 2020; Qiu et al., 2021), and for exploration of other neurobiological questions (Grydeland et al., 2016; Ma and Zhang 2017; Burt et al., 2018; Fukutomi et al., 2018; Li et al., 2019; Toschi et al., 2019; Gao et al., 2020; Liu et al., 2020; Paquola et al., 2020); however, additional considerations arise when using T1w/T2w myelin maps to address such questions. For example, although in the absence of head motion there is an exact correction of the spatially varying radiofrequency (RF) receive field (B1-) effects after taking the ratio of T1w and T2w images (either from the head coil or from the body coil after vendor receive field correction), this ratio leaves in RF transmit (B1+) field effects because they differ between the T1w and T2w images (Glasser and Van Essen 2011, and as will be discussed in Section 2.1 below). These B1+ effects are particularly noticeable in the HCP Young Adult (HCP-YA) dataset because of design compromises specific to that scanner^1^. These B1+ effects in the T1w/T2w myelin maps are most easily appreciated as correlated cross-hemisphere asymmetries in the cortical T1w/T2w ratio and B1+ maps (as will be noted in the results, e.g., Figure 5). Because such correlated asymmetries are not expected neurobiologically, are not present in alternative myelin mapping methods after B1+ correction (e.g., Carey et al., 2018), and were much smaller in the original Siemens 3T Trio datasets (Conte69 from Glasser and Van Essen 2011), an ad hoc correction method was initially developed to minimize them for HCP-YA data. This approach entailed subtracting the heavily surface-smoothed difference between a left/right symmetric group average Siemens 3T Trio T1w/T2w myelin map and each individual’s T1w/T2w myelin map, resulting in the “bias-corrected (BC)” MyelinMap (MyelinMap_BC) results produced by the HCP Pipelines and released by the HCP (Glasser et al., 2013; see Figure 18 therein).

Although this “MyelinMap_BC” approach works well for localizing cortical areas in individual participants, as the sharp transitions in cortical myelin content at areal boundaries are not affected by low spatial frequency B1+ transmit field biases (Glasser et al., 2016a), it was never intended for cross-participant statistical analyses of the sort mentioned above, as it removes both artifactual and genuine cross-participant T1w/T2w myelin map differences. Further, although the Siemens 3T Trio-based maps have reduced B1+ effects relative to the customized HCP-YA scanner, these remaining transmit effects are not zero, and are expected to be present on other scanner platforms as well. As a result, all prior T1w/T2w myelin maps contain at least some biases, as biases along the anterior-posterior and superior-inferior axes cannot be removed by symmetrization (such as a symmetric center versus periphery bias). Finally, the actual B1+ transmit field is modulated by the loading of the body coil by both the participant’s head and body, meaning that geometric variables such as head size and body size, and compositional variables such as weight and Body Mass Index (BMI), will have spurious correlations with uncorrected T1w/T2w myelin maps across participants. Such B1+ effects may correlate with other variables of interest, such as age or sex (MacLennan et al., 2021). To achieve the same flip angle in a particular location in the brain^2^, the scanner transmit reference voltage setting is calibrated higher for larger heads and bodies, compensating for the higher coil loading (as will be shown in Table 1). Further, the upper bodies of the participants induce asymmetries in the B1+ transmit field due to the circularly polarized transmit coil, and these asymmetries increase when the subjects bodies are larger (Glover et al., 1985; Sled and Pike 1998; Ibrahim et al., 2001; as will be shown in the results when comparing Figures 9 for children with smaller bodies on average and 12 for adults with larger ones obtained using the same sequences). These effects will occur across scanner vendors if circularly polarized body transmit coils are used.

**Table 1.** contains the Pearson correlations of relevant variables from the three human studies, HCP-YA (n=1042), HCD (n=628), and HCA (n=722).

| Table 1 | Sex |  |  | Age |  |  | BMI |  |  | HeadSize |  |  | Voltage |  |  |
| --- | --- | --- | --- | --- | --- | --- | --- | --- | --- | --- | --- | --- | --- | --- | --- |
| Study | HCP-YA | HCD | HCA | HCP-YA | HCD | HCA | HCP-YA | HCD | HCA | HCP-YA | HCD | HCA | HCP-YA | HCD | HCA |
| Sex | 1.00 | 1.00 | 1.00 | -0.22 | 0.04 | 0.02 | 0.06 | -0.04 | 0.03 | 0.73 | 0.45 | 0.67 | 0.55 | 0.26 | 0.39 |
| Age | -0.22 | 0.04 | 0.02 | 1.00 | 1.00 | 1.00 | 0.07 | 0.50 | -0.14 | -0.12 | 0.57 | 0.00 | -0.06 | 0.48 | -0.08 |
| BMI | 0.06 | -0.04 | 0.03 | 0.07 | 0.50 | -0.14 | 1.00 | 1.00 | 1.00 | 0.43 | 0.61 | 0.43 | 0.70 | 0.79 | 0.72 |
| HeadSize | 0.73 | 0.45 | 0.67 | -0.12 | 0.57 | 0.00 | 0.43 | 0.61 | 0.43 | 1.00 | 1.00 | 1.00 | 0.78 | 0.75 | 0.69 |
| Voltage | 0.55 | 0.26 | 0.39 | -0.06 | 0.48 | -0.08 | 0.70 | 0.79 | 0.72 | 0.78 | 0.75 | 0.69 | 1.00 | 1.00 | 1.00 |

Because the HCP consortium was aware of these limitations, an Actual Flip angle Imaging (AFI) scan (Yarnykh 2007), which measures the effect of the non-uniform B1+ transmit field on the flip angle obtained in every voxel, was acquired in most of the HCP-YA participants during the same imaging session as the T1w and T2w structural images. However, use of these AFI scans to improve the T1w/T2w myelin maps has remained unexplored until now. Additionally, subsequent HCP projects including the Human Connectome Development project (HCD) and Human Connectome Aging project (HCA) required shorter protocols than the HCP-YA project, necessitating removal of the AFI scans and creation of an alternative approach to B1+ transmit field mapping based on the images that were acquired. Finally, we apply the B1+ correction of T1w/T2w myelin maps to a non-human primate study, the Non-Human Primate Neuroimaging and Neuroanatomy Project (NHP_NNP), as T1w/T2w myelin maps are useful in cross-species comparisons (Glasser et al., 2014; Mars et al., 2018; Autio et al., 2020; Hayashi et al., 2021).

Here, we demonstrate a novel set of B1+ transmit field correction methods of T1w/T2w myelin maps using actual and “pseudo” measures of the B1+ transmit field (i.e., enabling correction even when a B1+ map was not obtained). We fit corrections of the T1w/T2w myelin maps at the group level, relying on (1) the hemispheric asymmetry of B1+ transmit field effects, and (2) the neuroanatomically grounded assumption that asymmetries in cortical myelin content should not be correlated with asymmetries in the B1+ transmit field at the group level to bootstrap corrected reference templates for a given species. We show that this correction removes not only the hemispheric asymmetries, but also additional symmetric biases. We then use these reference templates to correct individuals T1w/T2w myelin maps for B1+ transmit field effects without explicitly relying on hemispheric asymmetries. We show that this correction reduces B1+ correlated hemispheric asymmetries, symmetric biases, and spurious results in statistical comparisons of basic biological parameters of interest such as age, sex, or BMI across three independent human datasets. Finally, we compare the transmit and pseudo transmit correction methods in the more challenging macaque dataset where the effect is much smaller and closer to the noise floor due to a smaller brain size. The methods and results are organized as follows: Methods section 2.1 discusses the theory underlying the T1w/T2w approach and our new correction methods; Methods sections 2.2-2.6 present the four datasets, HCP-YA, HCD, HCA, and NHP_NNP, used in this study and the general preprocessing methods; Methods sections 2.7 and 2.8 respectively discuss the implementation of the real and pseudo-transmit correction methods, including specific issues pertaining to each dataset’s correction; Methods section 2.9 discusses specific issues related to transmit and pseudo-transmit field correction of macaques; Methods sections 2.10-2.13 address covariates for statistical models, “toy” statistical models illustrating the effects of B1+ on the relationship between sex, age, and BMI and T1w/T2w myelin, regressing covariates out of the maps to make improved maps for neuroanatomical use cases, and applying the correction in the volume (given that it is estimated on the surface). Results sections 3.1-3.3 present the results of transmit correction for HCP-YA, 3.4-3.5 present pseudo-transmit correction for HCD, 3.6-3.7 present pseudo-transmit correction for HCA, and 3.8-3.9 present both transmit and pseudo-transmit for NHP_NNP. Results section 3.10 addresses issues related to comparison between the transmit and pseudo-transmit approaches and test-retest issues.

## 2. Methods

### 2.1. Theory

#### The T1w/T2w approach to myelin mapping is an empirical approach that relies on several approximations

The first is that the ratio of the T1w and T2w images represent a combination of contrast of interest that is correlated with myelin content and a multiplicative bias field (Equation #1). In a given voxel, x^(3nd footnote)^ represents contrast for myelin and b represents the image intensity bias field (Glasser and Van Essen 2011).

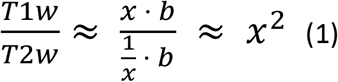

Ideally, the multiplicative bias field inherent in MRI images would cancel, leaving a bias free ratio image with increased contrast of interest. This ratio approach to bias correction has an advantage over algorithmic bias corrections in that it makes no assumptions about tissue intensity uniformity. There is a large literature exploring non-uniformity of the cortical grey matter—reviewed in Glasser et al., 2014—and the white matter also contains genuine non-uniformities (as will be shown in the results below, e.g., Figure 20). Importantly, these grey and white matter tissue non-uniformities vary independently, meaning that an algorithmic correction attempting to achieve white matter uniformity will induce a bias in grey matter and vice versa, resulting in a critical limitation of approaches that rely on algorithmic bias corrections using single tissues in isolation (e.g., Nerland et al., 2021). Indeed, this observation originally led to development of the T1w/T2w approach (Glasser and Van Essen 2011), because it makes no such assumption of tissue uniformity. Equation #1 can be expanded to approximate the effects of the receive (rb) and transmit (tb) bias fields separately (Equation #2)^4^.

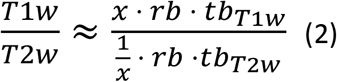

This simplifies to Equation #3, as the spatially inhomogeneous receive bias field cancels out exactly in the absence of motion and the myelin contrast is enhanced^5^.

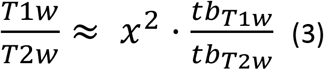

Unfortunately, tb_T1w_ / tb_T2w_ does not equal 1 in any location where the scanner does not achieve the prescribed flip angle, as T1w^6^ and T2w images experience different transmit field effects (see footnotes). Notably, the effect of tb_T1w_ / tb_T2w_ on T1w/T2w varies spatially according to the transmit field and is approximately multiplicative (Bonny et al., 1998; Collewet et al., 2002; Wang et al., 2004; Wang et al., 2005; Weiskopf et al 2011; Delgado et al., 2020) but does not substantially affect image contrast over the range of flip angles at 3T (Mugler 2014).

Importantly, these transmit field effects are expected to vary across participants (MacLennan et al., 2021) based on interindividual differences such as weight, Body Mass Index (BMI), or head size due to differential transmit coil loading and dielectric effects. These variables may themselves correlate with variables of interest such as sex or age, or even be intrinsically of interest themselves (such as BMI). Thus, if we have a measure of the transmit field (TF), we can use it empirically to attenuate these effects in the T1w/T2w data by determining the slope and intercept that relate the TF to the remaining bias in the T1w/T2w ratio with a linear multiplicative approximation, as in Equation #4 where tb_T1w_ / tb_T2w_ = 1/(TF*slope+intercept).

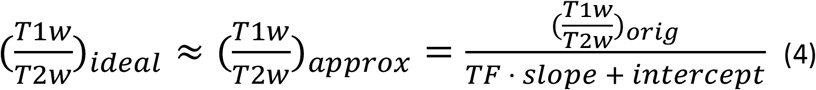

For simplicity, if the TF is already scaled such that a value of 1 means that the reference flip angle was reached exactly^7^ and we accept that the T1w/T2w values should not be changed where the achieved flip angle equals the nominal or reference value, the intercept must equal one minus the slope. This simplification leaves slope as the only parameter to optimize, resulting in Equation #5, which empirically is a good approximation (T1w/T2w)_approx_ for the much more complicated ideal (T1w/T2w)_ideal_ correction for the assumptions detailed below and in the footnotes above.

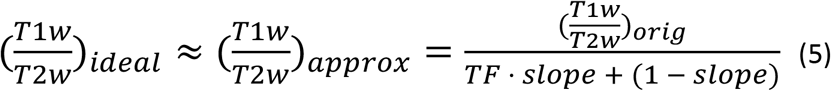

#### To optimize for the slope value of the B1+ transmit field correction, we created two different cost functions

For creating species-specific reference templates using left-right asymmetry, we use Eq. 6. For correcting individual subjects using a previously created species-specific reference template, we use Eq. 7. For the individual participant correction, comparison to a reference template is more robust because it does not require significant and consistent left-right asymmetry in the B1+ maps, which is variably present across individuals. Instead, Eq. 7 relies primarily on the highly consistent strong central to peripheral gradient in the transmit field that will be illustrated in the results, e.g., Figure 5, to fit the correction. To use Eq. 7, however, we first must have generated a corrected group reference template for that species (Eq. 6 is also used to find the value equivalent to the nominal or reference flip angle when using the pseudo-transmit approach described in section 2.8). We thus first create the reference template using the asymmetry method (Eq. 6). For Eq. 6 only, we make the plausible assumption that the correlated hemispheric asymmetries present in both the group T1w/T2w ratio and the group B1+ transmit field are spurious, as there is no neurobiological mechanism to generate such correlated asymmetries (nor are they seen in alternative non-invasive myelin mapping approaches with B1+ correction, including quantitative T1, T2*, and MT; Carey et al., 2018). This bootstrapping approach to generate a species-specific reference T1w/T2w template relies critically on the asymmetry of the B1+ transmit field typically found in 3T MRI scanners at the group level^8^, and would not have been possible with a symmetric B1+ transmit field. Thus, we used the cost function in Equation #6 to find the optimum slope to produce a corrected group T1w/T2w myelin map, where L is the left hemisphere and R is for the right hemisphere.

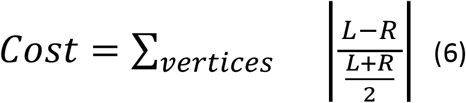

We chose to optimize this cost function using the golden search algorithm (Kiefer 1953; aka golden section search^9^). This optimization produces a group average transmit field corrected T1w/T2w myelin map (that we will present in the Results in Figure 6). Using this corrected T1w/T2w group surface map as a reference template (T), one can correct individual participants (I) using the more robust cost function in Equation #7.

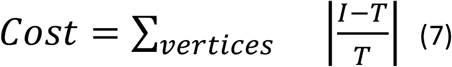

Importantly, when applying Equation #7, we want to allow for the existence of genuine per-participant differences in the spatial mean of the T1w/T2w ratio. To prevent such per-participant differences from biasing the cost function in Equation #7, we do the following: (1) identify the surface vertices where the calculated actual flip angle in the individual is close to the target flip angle (+/- 5%, where the B1+ correction will not cause appreciable changes to the values at those vertices); (2) find the median T1w/T2w ratio within those vertices for both the group and individual T1w/T2w maps; and (3) divide the ratio of the medians out of the individual map before computing the cost function. This process avoids over- or under-correcting the spatial pattern because of per-participant differences in the spatial mean of the T1w/T2w ratio (correction of nuisance effects on the T1w/T2w mean will be addressed in Methods section 2.10).

We demonstrate below that this approach indeed works if an AFI map is available, as is the case for the HCP-YA data (though unfortunately due to acquisition artifacts, the map is imperfect at the individual participant level, as will be described in Methods section 2.7 and will be shown in Figure 4). However, in some HCP-style studies no AFI map is available, so an alternative approach is needed. A candidate alternative emerged in the course of developing an improved receive field bias correction approach for fMRI data termed the “SEBASED” correction (Glasser et al., 2016a), which enables biological interpretation of fMRI parameter maps (e.g., so that estimated task fMRI betas or variances have a consistent scale across space). In particular, differential transmit field effects were also present in the gradient echo EPI (echo-planar imaging) fMRI images and the spin echo EPI field map images [for the same reason that they are present in the T1w and T2w images as described above, i.e., the signal intensity depends on sin(FA*TF) for gradient echo images and sin^3^(FA*TF) for spin echo images, FA=Flip angle; TF=Transmit Field; Bonny et al., 1998; Collewet et al., 2002; Wang et al., 2004; Wang et al., 2005; Weiskopf et al 2011; Delgado et al., 2020]. For the purposes of fMRI receive bias correction, such transmit effects were nuisances that needed to be appropriately modeled (together with T2* induced susceptibility dropouts) to generate an accurate receive field estimate. However, this “pseudo” transmit field can also be used for T1w/T2w myelin map correction if it is collected during the same session as the structural data. Indeed, at the group average level, this pseudo-transmit field has a remarkably similar spatial pattern to the AFI-based transmit field (as will be shown when comparing Figures 5 versus 9 and 12). Thus, the pseudo-transmit field can be used in the same way as a real transmit field if a reference value can be established, analogous to the nominal or reference flip angle in the AFI transmit field (as will be described in the pseudo-transmit specific methods section 2.8 below).

#### The above theoretical considerations entail several assumptions

1) *No significant change in head position between the T1w and T2w images*. For the receive field to exactly cancel, it must be identical in both images. Because the head coil is stationary during a scan, head motion changes where the brain anatomy is inside the receive bias field. The use of on-scanner corrections such as Siemens PreScan Normalize will mitigate this issue, as a fixed, sharply varying receive field from the head coil is removed from the images by the scanner, and thus prior to downstream motion correction (which will align the anatomy but not the receive fields, given the differential relationship between brain anatomy and B1-receive field when head motion occurs). Thus, if on-scanner B1-corrections are not used, it is helpful to separately acquire data to measure the B1-receive field and apply this correction to the data in a way that accounts for the head movement within a static bias field (as will be described in methods section 2.7). This assumption also applies to the use of GRE and SE EPI images for pseudo-transmit field computation, as will be described in methods section 2.8.

2) *No significant change in head position between the B1+ transmit field map and T1w and T2w images*. Changes in head position may intrinsically change B1+ inhomogeneity due to differential coil loading and reduce the accuracy of B1+ field correction. This assumption is difficult to address, but it is expected to be a small effect at 3T given most 3T scanners use a large body transmitter coil and thus that the B1+ field will be substantially smoother than the B1-field. This is also not an issue for non-human primate imaging because the head is almost always fixed during imaging.

3) *The B1+ transmit field can be related to the B1+ bias in the T1w/T2w images with a linear multiplicative approximation (Eq. 5)*. This manuscript provides empirical support for these assumptions for the field strength and sequences used here, but, as we note above in the footnotes and in the discussion, an exact correction would be much more complicated. Our success here is likely the collective result of optimized hardware conditions, chosen sequence parameters, and well-implemented key features of each pulse sequence (e.g., the adiabatic inversion-recovery pulse in the MPRAGE (Garwood and Uğurbil 1992) and a SPACE sequence where the typical non-linear effect of the B1+ field on spin echo-based T2w images is conditioned by an optimized variable flip angle train that yields a small and approximately linear dependence of signal intensity and contrast on flip angle variation (i.e., ∼+/-20% typically observed in the human brain at 3T (Mugler 2014)). Application of this method outside of these parameters, e.g., at higher fields such as 7T or using different pulse sequences would benefit from further validation to ensure that the kinds of B1+ effects that we attenuate here are still being removed.

4) *The pseudo-transmit field reference value correctly indicates where the scanner achieves the intended flip angle.* Individual differences in T2 and T2* may influence the pseudo-transmit field map beyond the real transmit field. T2* from large scale magnetic field inhomogeneity is dealt with using the thresholding described below, but it is clear that the estimated reference value is slightly different in the HCD and HCA datasets due to such effects. Similarly, the scanner is assumed not to differentially scale the GRE and SE images automatically in different subjects (and such behavior has not been found on Siemens scanners).

### 2.2. Participants

Data from the Human Connectome Young Adult (HCP-YA), Development (HCD), and Aging (HCA) Projects were acquired and used with approval of the Washington University Institutional Review Board (IRB) and those of partner institutions (Van Essen et al., 2013; Bookheimer et al., 2019; Harms et al., 2018; Somerville et al., 2018). The Non-Human Primate Neuroimaging and Neuroanatomy Project (NHP_NNP) data were acquired with approval from the RIKEN Kobe Japan institutional animal care and use committee (Autio et al., 2020; Hayashi et al., 2021). Figure 1 illustrates the cohorts, specific analyses, and corrected T1w/T2w outputs produced in this study. For HCP-YA, 1042 participants had successful MSMAll registration and had AFI scans available (ages 22-35, 470 men and 572 women, BMI 16.5-47.8, head size 2433cc to 4461cc, and body coil reference voltage 183V-356V). For HCD, 628 participants had successfully completed preprocessing and quality control at the time of the study and were publicly released as part of the “Lifespan HCP Release 2.0” [ages 8-22 (5 to 7-year-olds were not included due to small numbers at this time), 292 boys and 336 girls, BMI 13.1-43.2, head size 2133cc to 4260cc, and reference voltage 206V to 317V]. For HCA, 722 participants had successfully completed preprocessing and quality control at the time of the study and were publicly released as a part of the “Lifespan HCP Release 2.0” (ages 36-90+, 319 men and 403 women, BMI 15.7-43.7, head size 2522cc to 4719cc, and reference voltage 217V-337V). The correlations of these demographic variables are listed in Table 1. The NHP_NNP acquired data from 16 macaque monkeys (13 *M. mulatta* and 3 *M. fasicularis*, 16 male and 0 female, ages 3.5-7.0, weights 3.5-9.8kg, head size 342cc to 570cc, and reference voltages 266V-282V). For all studies, a consistent imaging protocol was used within the study, enabling comparisons across subjects within each study. The data for this study are available at the BALSA neuroimaging study results database (https://balsa.wustl.edu/study/show/mDBP0; Van Essen et al., 2017), and a link to each figure’s specific data is provided in the legend. Note that the figures can be exactly replicated on the reader’s computer by downloading and loading the appropriate ‘scene’ file in Connectome Workbench, thus providing a convenient mechanism to visually explore the results in greater detail or with different color palette scaling. The methods presented in this study will be released as an HCP Pipeline (https://github.com/Washington-University/HCPpipelines).

**Figure 1.**
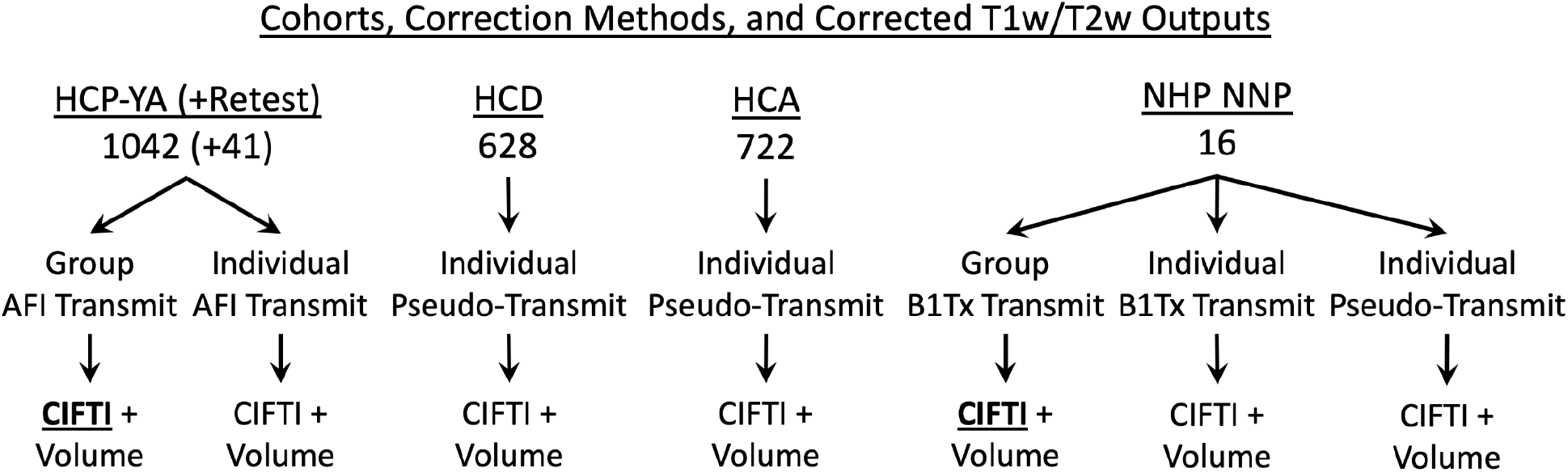
illustrates the datasets, specific analyses, and corrected outputs in this study. **<u>Bold</u> <u>underlined</u>** font indicates the reference T1w/T2w outputs (which are computed only for the surface). Surface data are stored in the CIFTI format (Glasser et al., 2013), which combines both hemispheres into a single file.

### 2.3. Scans HCP-YA

The acquisition of the HCP-YA data has been described previously (Glasser et al., 2013; Uğurbil et al., 2013). Briefly, 0.7mm isotropic resolution T1w MPRAGE images (Mugler and Brookeman 1990) were acquired with TI=1000ms, flip angle=8 degrees, TE=2.14ms, and TR=2400ms. 0.7mm isotropic resolution T2w SPACE images (Mugler et al., 2000) were acquired with TE=565ms and TR=3200ms. One or two T1w and T2w images were used depending upon image quality. A 4.5 min AFI scan (Yarnykh 2007) was acquired at 2mm isotropic resolution, TR1=20ms, TR2=120ms, 50-degree nominal (i.e., reference) flip angle, and TE=1.9ms. Additionally, 2mm isotropic “Bias_32CH” and “Bias_32BC” images were acquired with a gradient echo sequence (TR=250ms, TE=1.01ms) with the only difference being whether the 32-channel head coil (32CH) or body coil (BC) was used for image receive (each ∼ 0.5 min). These images were acquired to measure the B1-receive field, as it was desired in the HCP-YA to do all image processing offline and Siemens PreScan Normalize (PSN) was not used for any acquisitions. (For simplicity and robustness, however, we recommend the use of Siemens PreScan normalize for future studies for all acquired images including structural, functional, diffusion, and spin echo field maps). The images were obtained on a customized Siemens 3T ‘ConnectomS’ scanner with 100mT/m gradients using a Siemens standard 32 channel head coil. All acquisitions used the body coil for radiofrequency transmission. All images had the face and ears removed using a defacing algorithm (Milchenko and Marcus 2013) to prevent identification of participants. Forty-one Test-Retest participants were scanned and processed a second time through all modalities (interval: 18-328 days; mean: 135 days). In a separate scanning session, gradient echo EPI fMRI data were acquired at 2mm isotropic resolution along LR and RL phase encoding directions with matching phase reversed spin echo fMRI data (Smith et al., 2013), data which were used for MSMAll areal feature-based surface alignment (as will be described in section 2.6 below).

### 2.4. Scans HCA and HCD (HCP-Lifespan)

The acquisition of the HCD and HCA data has been described previously (Harms et al., 2018). Briefly, 0.8mm isotropic resolution T1w multi-echo MPRAGE images (van der Kouwe et al., 2008) were acquired with an internal EPI-based navigator (Tisdall et al., 2012; vNav) with TI=1000ms, flip angle=8 degrees, TE=1.8/3.6/5.4/7.2ms, and TR=2500ms. Only the average of the first two echo times were used because of artifacts that were subsequently discovered in the later echo times (a multi-echo MPRAGE was used in an effort to equalize receiver bandwidth between the T1w and T2w images (van der Kouwe et al., 2008); however, such an approach is not required as readout distortions can be corrected in T1w and T2w images (Glasser et al., 2013), and a short TE single echo MPRAGE as used in the HCP-YA has higher SNR and fewer artifacts; Elam et al., 2021). 0.8mm isotropic resolution T2w SPACE images (Mugler et al., 2000) were acquired with an internal EPI-based navigator (Tisdall et al., 2012; vNav) with TE=564ms, TR=3200ms. Spin echo (SE) EPI images with both anterior-to-posterior (AP) and posterior-to-anterior (PA) phase encoding polarities were acquired at 2mm isotropic resolution. Gradient echo (GRE) EPI-BOLD fMRI images were acquired with identical geometric specifications, again with both AP and PA polarity. These images were obtained on a standard Siemens 3T Prisma with 80mT/m gradients using a Siemens standard 32 channel head coil. For the T1w and T2w acquisitions, both PSN and unfiltered (non-normalized) reconstructions were generated. The unfiltered versions were used in initial HCP Pipelines preprocessing to match the approach used in HCP-YA, though this is no longer recommended (PSN is now recommended for simplicity and robustness). Defacing (Milchenko and Marcus 2013) was run on the 3D T1w and T2w images including separately on the PreScan normalized and unfiltered images.

### 2.5. Scans NHP_NNP

The acquisition of the NHP_NNP data has been described previously (Autio et al., 2020). Briefly, 0.5mm isotropic resolution T1w MPRAGE images were acquired with TI=900ms, flip angle=8 degrees, TE=2.2ms, and TR=2200ms. 0.5mm isotropic T2w SPACE images were acquired with TE=562ms and TR=3200ms. B1+ mapping (“B1Tx”) was acquired using a vendor-provided slice-selective RF-prepared sequence (called ‘B1map’ by Siemens) with a turbo fast low-angle-shot (FLASH) read-out with 2.0mm isotropic resolution, flip angle=8 degrees, nominal target flip angle=80 degrees, TE=2.5ms, and TR=20s (Chung et al., 2010). Because the preparation RF pulse excites slightly thicker slices than the FLASH acquisition (requiring a gap between the slices), even and odd slices (with a gap 2 mm) were acquired in separate runs for full brain coverage (each 40s in duration). A pair of reversed phase-encoding (left-to-right [LR] and right-to-left [RL]) spin echo EPI images were acquired at 1.25mm isotropic resolution together with reversed phase-encoding gradient echo EPI-BOLD fMRI images with identical geometric specifications. These images were obtained on a standard Siemens 3T Prisma with 80mT/m gradients using a customized macaque-sized 24 channel head coil (Autio et al., 2020) that enables HCP-style pulse sequences to be acquired in the macaque (commercially available from Rogue Research Inc., Montreal/Takashima Seisakusho Co., Tokyo). PSN was used on all images (as recommended for all human and non-human primate studies going forward for simplicity and robustness). The macaques were anesthetized using dexmedetomidine and low-dose isoflurane in a protocol described previously (Autio et al., 2020).

### 2.6. General Preprocessing

All datasets were preprocessed using the HCP Pipelines (Glasser et al., 2013) or HCP-NHP Pipelines (Autio et al., 2020), which included the structural spatial preprocessing pipelines, the functional preprocessing pipelines, the multi-run spatial ICA+FIX (Salimi-Khorshidi et al., 2014; Glasser et al., 2018) fMRI data denoising pipelines, and, in humans, multi-modal ‘MSMAll’ areal-feature-based surface registration (Robinson et al., 2014; 2018; the utility of MSMAll has yet to be established in the less variable macaque brain so cortical-folding-based ‘MSMSulc’ registration was used in macaques). The relevant results of these pipelines for the purposes of this study are MSMAll (or MSMSulc in the case of macaques) aligned cortical T1w/T2w myelin maps and also T1w/T2w ratio volumes in MNI space (aligned using nonlinear FSL FNIRT-based registration of the T1w image). Figure 2 illustrates the general T1w/T2w transmit or pseudo-transmit correction pipeline operations and outputs with multiple optional steps indicated. This pipeline will be described in detail over sections 2.7 and 2.8. The group transmit (for generating reference T1w/T2w myelin maps), group pseudo-transmit (for generating reference values) and individual transmit and pseudo-transmit nested optimization algorithms are illustrated in Figure 3 and will be described in sections 2.7 and 2.8 below.

**Figure 2.**
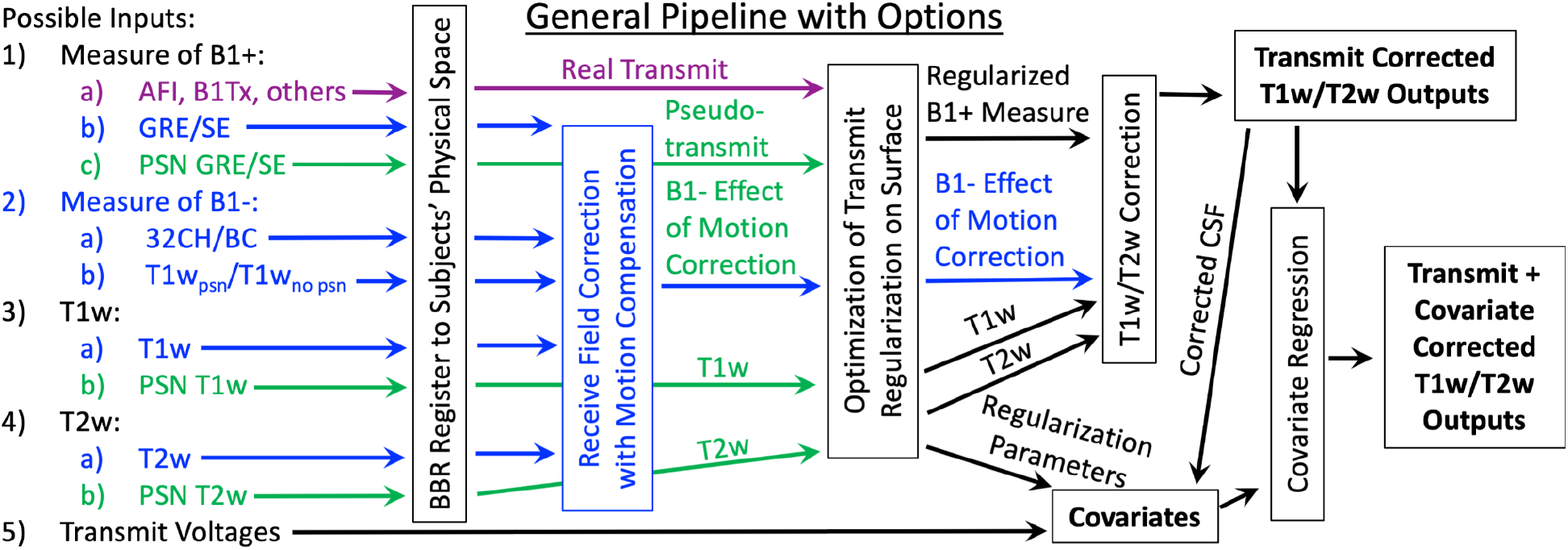
illustrates the overall T1w/T2w transmit field correction pipeline operations and outputs, with inputs and outputs represented by horizontal boxes and processing steps by vertical boxes. Options (depending on specifics of data acquisition) are indicated with different colors (purple for real transmit field, green if Siemens PreScan Normalize (PSN) was used on all images, and blue if motion compensation for the receive field is needed because PSN was not used). Lines that intersect a step (without an arrowhead/passing under the step) skip that particular step. The simplest form of the pipeline (green path) would require that the T1w and T2w images (and GRE/SE images if the pseudo-transmit approach is used) have undergone PSN or the equivalent from another vendor, which permits skipping the Receive Field Correction with Motion Compensation step (blue paths). **Bold** text indicates the primary outputs of interest for downstream analyses. In all cases, outputs are in CIFTI, NIFTI physical volume, and NIFTI MNI volume spaces.

**Figure 3.**
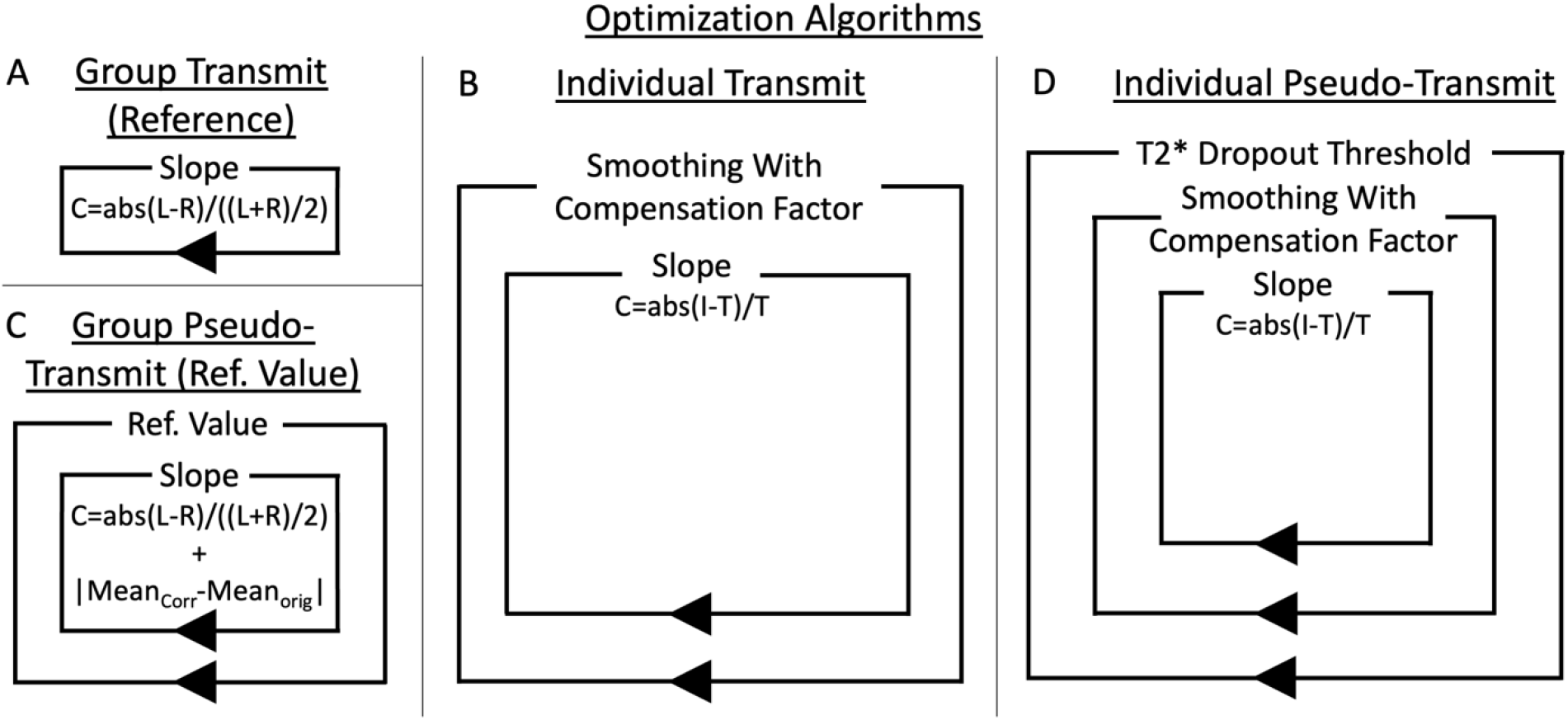
illustrates the four optimization approaches used in this study. Each box represents a loop, and anything the box surrounds is inside that loop body. The simplest is the group transmit optimization approach (A), which uses a single loop with the cost function in Equation #6 and generates species specific reference T1w/T2w myelin maps (the group average B1+ transmit fields are assumed not to require regularization due to cross-participant averaging for removal of artifacts). The reference map is then used in the individual transmit nested optimization algorithm (B) together with the cost function in Equation #7. The outer loop finds the optimal amount of (spatially constrained) smoothing (for regularization) while compensating for the change in fraction of flip angles above and below the reference flip angle (compensation factor) and the inner loop finds the optimal slope. For the pseudo-transmit approach, the appropriate reference value (the GRE/SE value where the target flip angle is expected to have been achieved by the scanner) must also be found for a given study together with the slope, which is again done at the group level using Equation #6 with the addition to the cost function of the absolute value of the change in the overall group mean (C). Then the individual pseudo-transmit nested optimization algorithm (D) is used together with the cost function in Equation #7, with the addition, relative to the individual transmit algorithm (B), of a T2* dropout threshold optimization loop. The reference map from the group transmit algorithm is used in both transmit and pseudo-transmit approaches (because the unregularized pseudo-transmit maps used in (B) have regions with substantial T2* dropout). Appendix A contains a pseudocode representation of the individual pseudo-transmit algorithm.

### 2.7. AFI Image Preprocessing

The raw AFI volumes consist of two image volumes acquired at two different TRs. These need to be corrected for gradient nonlinearity distortions (as the other images have been) and rigidly aligned to the T1w scans in the participant’s physical space. FreeSurfer boundary-based registration (BBR) (Greve and Fischl 2009) is used for this alignment after initialization with 6 DOF FSL FLIRT registration. At this point, the raw AFI volumes are aligned to the participant’s physical space (and can be trivially resampled to MNI space or mapped onto the cortical surfaces). Unfortunately, the second AFI volume invariably contains irregular ringing artifacts, which induce artifacts in the computed flip angle maps (Figure 4; Yarnykh 2007). These artifacts are most likely due to inadequate spoiling and crusher gradients for the second volume (Nehrke 2009), but were of variable severity across the HCP-YA participants. We opted to simply average across participants to eliminate them at the group level for the purpose of this manuscript.

**Figure 4.**
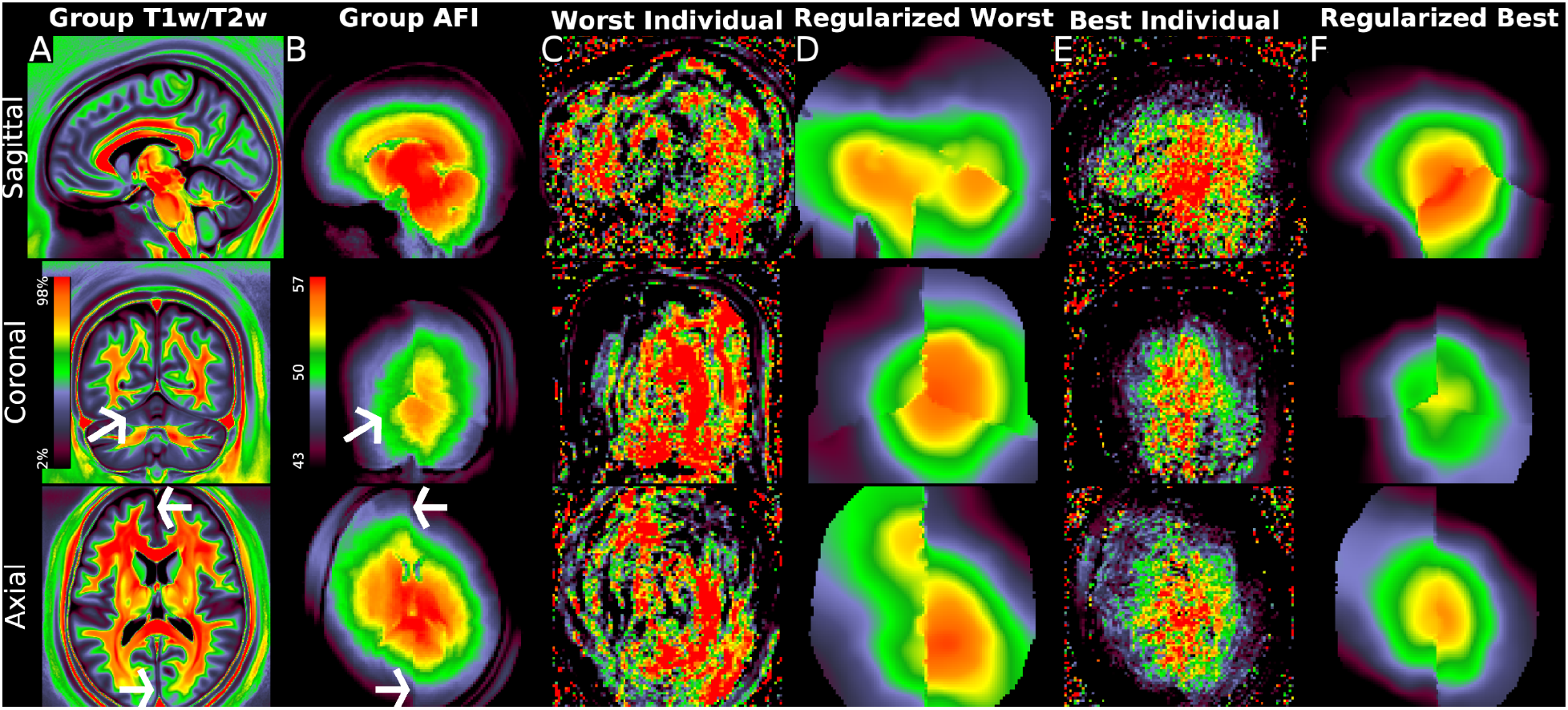
illustrates the group average original T1w/T2w volume and the unsmoothed group average AFI map together with unregularized and regularized AFI maps from two exemplar individuals. Panel A shows the group average original T1w/T2w average volume in sagittal (top), coronal (middle) and axial (bottom) slices. Panel B illustrates the group average AFI map, where the original AFI volumes (i.e., TR1 and TR2) are aligned to MNI space nonlinearly and then averaged across participants without smoothing; the AFI flip angle map is then computed from these results at the group level. The arrows mark the anterior and posterior falx and the left tentorium, which all show sharp discontinuities in the AFI map, indicating that the B1+ field is particularly affected by these fibrous dural reflections. Panels C and D illustrate a participant with the worst (worst correlation with the group average map) ringing artifacts before and after regularization. Panel E and F illustrate a participant with the least (best correlation with the group average map) ringing artifacts before and after regularization. Other participants lie between these extremes. Additionally, Panel A shows a subtle correlation with the hemispherically asymmetric pattern in the B1+ map in Panel B within the deep white matter; however, contrast differences between tissues such as CSF/grey matter/white matter are much stronger than those within tissues and also the B1+ effect. This map also illustrates how partial volume effects between CSF/grey matter/white matter tissues overwhelm differences in myelin content within the cortical grey matter in group average T1w/T2w volume maps, making them inappropriate for cortical analyses. Methods sections 2.2 and 2.3 describe the participants and data and preprocessing is described in methods sections 2.6 and 2.7. https://balsa.wustl.edu/PrBrK.

Although HCP-style studies always avoid 3D spatial smoothing to preserve spatial localization ability for structural and functional imaging analyses (because most brain neuroanatomy is not spatially smooth in 3D; Glasser et al., 2016b; Coalson et al., 2018), here we can use the property that the B1+ transmit field maps are (mostly) smooth in 3D to reduce the artifacts in individual subjects in the AFI maps. That said, Figure 4 illustrates sharp discontinuities in the 3D structure of the B1+ transmit field at the fibrous falx and the tentorium in group average data without smoothing. This interesting property of the B1+ transmit field is presumably related to the electrical properties of different brain tissues (Vaidya et al., 2016), but has not, to our knowledge, been previously shown at this level of anatomical detail, and it has important implications for transmit field map regularization. When we initially used uniform smoothing in 3D, we found good correction of lateral cortex but residual bias medially, indicating that the fibrous dural tissue’s effect on B1+ is present (though not easy to see) in the T1w/T2w images themselves. Better results that eliminated the residual medial bias were obtained by smoothing using the same kernel FWHM within three distinct regions generated from FreeSurfer’s individual participant segmentations – the left and right cerebral hemispheres separated by the falx, and the combination of the cerebellum and brainstem separated by the tentorium. We did allow smoothing across the corpus callosum and the midbrain, as they lack a fibrous discontinuity (anatomically and in the transmit field). Because valid flip angle data is present outside the grey and white matter (Figure 4), we also allowed smoothing to occur within the rest of the entire head to reduce edge effects.

We smoothed in the participant’s physical space because that was where the B1+ was generated. Because the artifacts vary in severity on a per participant basis (Figure 4), we used a golden search approach (similar to what was used above for finding the slope with Equation #7) to optimize the amount of smoothing in FWHM in each participant (see Figure 3). Finally, to avoid introducing biases in the T1w/T2w myelin map mean from the smoothing because heavy smoothing moves the average location of the reference flip angle towards the central regions of higher flip angles (because the mean flip angle is less than the reference flip angle), we compensated the smoothed maps (multiplying by a factor) to ensure that a similar number of voxels are above and below the reference flip angle so that the average location of the reference flip angle is in the same location as it was prior to smoothing. At this point the final flip angle map in volume or surface space is ready for correction of the T1w/T2w myelin map using Equation #5.

The HCP-YA data have an additional consideration (relative to the above Theory section): they were not acquired with PreScan Normalize. Instead, two short gradient echo scans were acquired for estimating the receive bias field. These two scans (“BIAS_32CH” and “BIAS_BC”) differed only in the receiver coil used (32-channel head coil vs body coil). Such data were not used when running the HCP Pipelines originally and thus the original T1w/T2w myelin maps have biases in them related to participant motion between the T1w and T2w scans. Thus, we used these scans to generate a smoothed (8mm FWHM) receive field restricted to the brain (due to poor estimation in osseous structures and differential defacing effects in the two images) and then dilated with extrapolation to estimate the field outside the brain to enable correction even in the setting of bulk head motion between the receive field scans and the T1w and T2w scans. To align this data to the T1w image, we used FreeSurfer BBR registration (Greve and Fischl 2009) using the pial surface instead of the default white surface, given that grey matter/CSF contrast exceeded grey/white contrast in these images. This field was then applied so as to match the estimated participant movement between the T1w and T2w images (averaging the receive fields if two T1w or T2w images were used prior to dividing T1w by T2w) to produce a final field that represented the error from motion on the original T1w/T2w receive field correction. This “error in receive field correction” field was then used to correct the T1w/T2w ratio data prior to fitting the AFI transmit field map and final T1w/T2w transmit field correction.

### 2.8. Pseudo-transmit Field Generation and Preprocessing in Humans

As described above in section 2.1, the idea behind using gradient echo and spin echo EPI images to compute a “transmit field”-like measure is that, just as the gradient echo T1w MPRAGE and spin echo T2w SPACE sequences experience differential transmit radiofrequency effects, gradient echo fMRI and spin echo field map images also experience different transmit radiofrequency effects^10^. Unfortunately, they experience other effects that must also be considered, including magnetic inhomogeneity effects in the gradient echo image (i.e., signal dropout in regions of main magnetic field (b0) inhomogeneity from air-tissue interfaces, around blood vessels, and at tissue boundaries such as the brain and skull), potentially differential receive fields (if there is head motion between scans that did not all use PreScan Normalize), and slight differences in tissue contrast (due to T2* versus T2 weighting). These issues are addressed using the approaches described in the next two paragraphs. Additionally, magnetic field inhomogeneity causes both types of images to suffer from signal pile up and rarefaction, and gradient echo images additionally have their intensities modulated by differential effective echo times across space (Deichmann et al., 2002; De Panfilis and Schwarzbauer 2005), requiring correction using a combination of Jacobian modulation and combining across opposed phase encoding directions. We used balanced amounts of data from opposed phase encoding directions to ensure that such contributions are appropriately compensated.

To produce the pseudo-transmit field, the average across phase encoding polarities of the single band reference gradient echo images (“SBRef;” after alignment and distortion correction) is divided by the corresponding average from the Jacobian modulated spin echo images. This division cancels the receive field (see next paragraph) and markedly reduces the tissue contrast in the image while preserving differential B1+ effects. For the pseudo-transmit approach, it is necessary to determine the “reference value” of the pseudo-transmit field that represents the location where the scanner achieves the prescribed flip angles^11^. The reference value is found at the group level using the group average uncorrected myelin map and “raw” group average GRE/SE images, where first a reference value and then a slope are fit in a nested algorithm whose cost function minimizes the left to right asymmetry (Equation #6; see Figure 3). When estimating the reference value, we also penalize changes to the overall group mean to ensure a unique solution (as our simplification allowing representation of the intercept in terms of the slope in Equation 5 will not hold when fitting both the reference value and slope if the mean is allowed to change without penalty). Because the raw group average GRE/SE images have significant T2* dropout regions, we did not produce a reference map from this correction for HCD and HCA, but instead used the species-specific reference myelin map computed from the transmit field approach in the HCP-YA data or NHP data. Several remaining free parameters must be determined, which we accomplish in a nested fashion using the same golden search algorithm described above (i.e., the golden search for T2* dropout threshold (1) uses a cost function that runs an internal golden search on the smoothing (2), which in turn uses a cost function that internally runs a golden search on slope (3) as illustrated in Figure 3, and with pseudocode in Appendix A)^12^. *1) The threshold at which the GRE / SE-EPI pseudo-transmit field is contaminated with T2* dropout effects.* If the threshold is too small, holes in the field will appear in regions of susceptibility dropout, but if it is too large, the range of the field will be clipped for low flip angle regions. *2) The smoothing FWHM to reduce noise and tissue-specific effects* (using the same constrained 3D volumetric smoothing approach described above for the AFI scans with the same approach for compensating for the effects of smoothing on the mean of the pseudo-transmit field map). After thresholding but prior to smoothing, extrapolating dilation (using the local gradient to predict continued changes outside of the valid data region) is used to fill the head mask to avoid edge effects of smoothing at the edge of the valid data. This dilation is necessary because, unlike the AFI approach above, valid data is not present outside of the brain because of the application of fat saturation to the gradient echo and spin echo EPI acquisitions. The effect of the smoothing is to reduce tissue contrast differences from differential T2* and T2 weighting. *3) The slope of the T1w/T2w pseudo-transmit field correction* (to use in Equation #5). The slope is fit using Equation #7 as the cost function.

Additionally, for the HCP-Lifespan data the T1w and T2w images were acquired with PreScan Normalize, but the unnormalized versions were used for preprocessing for historical reasons (which is not recommended going forward for simplicity and robustness). Thus, the effect of any head motion between the T1w and T2w images (and GRE and SE images) must be compensated for, as described above. Explicit “BIAS” scans for estimating the receive field were not collected for HCP-Lifespan, however. Thus, for HCP-Lifespan, the receive field was computed by dividing the non-PreScan Normalized T1w image by the PreScan Normalized T1w image (the T2w image had additional on-scanner filters applied and could not be used in this way). Since these images entered the de-facing pipeline separately, de-facing differences were present due to the large discrepancies in image intensity and bias that affected the defacing pipeline registration algorithms (Milchenko and Marcus 2013). Thus, values outside of the brain for the receive field were unreliable and had to be inferred through extrapolating dilation as described above. Once computed, the effects of motion on this receive field can be modeled for the T1w and T2w images together with the GRE and SE images to ensure that the receive field properly cancels in all ratio images (T1w/T2w and GRE/SE).

### 2.9. Special Considerations for Macaque Transmit Field and pseudo-transmit Field Preprocessing

For the macaques, the data necessary for both real and pseudo-transmit field correction were available in the same scanning session, so the two approaches could be directly compared. The transmit field consists of a “magnitude” image and a “phase” image^13^. The magnitude image contains artifacts outside the brain that make registration initialization problematic. Because the macaques were anesthetized with their head fixed using adhesive medical tape, motion between the scans is expected to be minimal. Thus, registration was initialized using the existing FreeSurfer-based BBR registration between the scanner space fMRI and T1w images (under the assumption that the brain did not move much between the fMRI and B1Tx acquisitions) and then was fine-tuned with FreeSurfer BBR (Greve and Fischl 2009) to generate a precise registration between the B1Tx and T1w data. This approach avoids the need for failure-prone unmasked registration or arduous brain extraction of the B1Tx magnitude images for the initialization. Similar to the other measures, the transmit field has some artifacts that are reduced by falx and tentorium constrained volumetric smoothing. Given the finer spatial resolution available and smaller brain sizes, distances for extrapolating dilation and outlier detection were reduced by half for the macaque relative to human scans. Because Siemens PreScan Normalize was used on all images, post-hoc receive field correction for motion was unnecessary and was skipped (Figure 2). As above for HCP-YA, a macaque T1w/T2w reference myelin map was generated using the unregularized group average transmit field and T1w/T2w myelin maps with the L-R cost function (Equation #6), and then individual participants were corrected with both the transmit field and pseudo-transmit field approaches using the I-T cost function (Equation #7). Because pseudo-transmit and real transmit data were available in the same group of subjects, the reference value was computed exactly as the mode of the pseudo-transmit values in the locations whose transmit value was +/-1% of the nominal transmit value.

### 2.10. Covariates for Downstream Statistical Analyses Involving Groups of Individuals

The above methods aim to markedly reduce the spatially non-uniform effects of variable B1+ inhomogeneity, which correlate with interindividual differences such as BMI and head size and which may correlate with variables of interest in downstream statistical analyses. They also address the effects of motion on B1-receive field correction across participants which introduce random artifacts in T1w/T2w myelin maps. Nonetheless, the T1w/T2w ratio is not an intrinsically quantitative measure such as, for example, a tissue T1 value. Because it has the potential to vary, we produce reference T1w/T2w values from the lateral ventricular CSF, which are not expected to vary with myelin content, as a guard against any effects from the scanner that are not described in a consistently acquired protocol, or unexpected effects from the correction algorithm. Because there may be differences across ages in ventricle size and CSF flow artifacts, we chose the CSF T1w/T2w percentile (searching from 1^st^ to 99th percentile) whose values had the strongest correlation with the mean T1w/T2w value across participants (after eroding the lateral ventricles by 1 voxel to eliminate partial volume effects). Additional covariates we consider appropriate to include are measures of within-scan head motion if available (e.g., when using a 3D MPRAGE or SPACE with a navigator) and the correction parameters mentioned above (slope, mean B1+ measure, smoothing FWHM, pseudo-transmit threshold, and the compensation factor required to account for the effects of smoothing). Finally, we found that the reference voltage of the transmit coil was a useful additional covariate of no interest that helped to further remove dependence on the scanner generated B1+ field, given the correlation between the mean T1w/T2w and the transmit voltage (r=0.44 in the HCP-YA data). Overall, the purpose of the covariates is to attempt to account for nuisance variables that may affect the spatial mean of the T1w/T2w myelin maps. Such remaining nuisance effects on the spatial mean are possible because the transmit field correction intentionally avoids removing differences in the spatial mean between the individual and reference map, to avoid removing real neurobiological differences (see paragraph in methods section 2.1 immediately following Equation #7 above). For example, the correction of the spatially non-uniform effects of the B1+ field only modestly decreases the correlation between mean T1w/T2w and the transmit voltage (r=0.37 in the HCP-YA data). Thus, the covariates correct the B1+ effects on the mean T1w/T2w.

### 2.11. Toy Statistical Model for Demonstrating the Effects of Individual participant T1w/T2w Myelin Map Correction

To illustrate the beneficial effects of T1w/T2w myelin map correction on cross-participant analyses that relate participant demographics, behavior, and disease states to T1w/T2w myelin maps, we developed a “toy” statistical model. The variables of interest are sex, age, and Body Mass Index (BMI). We use multiple linear regression after normalizing the input variables to demonstrate simple linear relationships within the 360 cortical areas identified in the Human Connectome Project’s multi-modal parcellation V1.0 (Glasser et al., 2016a). Importantly, in this work we focus exclusively on the effects of bias and nuisance from spurious B1+ variations on the T1w/T2w myelin maps used in these linear regressions and not on any neuroscientific questions of interest. Such neuroscientific questions will be properly addressed in subsequent publications using more advanced statistical models (e.g., Baum et al., 2021 for development). For this reason, we do not assess the statistical significance of these effects and will comment on them only qualitatively insofar as it is useful to describe the effects of the transmit field correction.

### 2.12. T1w/T2w Myelin Maps for Neuroanatomical Use Cases After Transmit Field Correction

Although the covariates can be included in statistical models as described above, it is also helpful to produce T1w/T2w myelin maps with these parameters regressed out of the data (to correct the individual participant myelin map spatial means) for neuroanatomical use cases, such as parcellation. Thus, we also produced T1w/T2w myelin maps with these parameters regressed out, which will be included in the boxplot figures in the results section 2.10 below (Figures 18 and 19). Despite all these corrections, these maps may have some residual artifacts, for example, from (1) within-scan head motion (either in scans without the vNav motion correction or where the limit on the number of additional TRs reacquired due to motion is met), or (2) head coil malfunction in one scan but not the other (T1w or T2w). Thus, the “MyelinMap_BC” approach (described in the Introduction and in Glasser et al., 2013), but using the new B1+ corrected reference group myelin map, may still be preferable to these regressed T1w/T2w myelin maps in such situations for neuroanatomical use cases, given that even covariate regression cannot eliminate all forms of low spatial frequency myelin map biases that may occasionally occur.

### 2.13. Correction of T1w/T2w Ratio Volumes

Although the relationship between the T1w/T2w ratio and myelin is less well constrained outside of cortical grey matter (Glasser and Van Essen 2011), some investigators have been interested in exploring the T1w/T2w ratio in subcortical structures, including white matter. For all above data, the B1- corrections were applied to the T1w/T2w volume appropriately, and the B1+ transmit field measure was regularized initially in volume space before being projected to the surface as described above in sections 2.7 and 2.8. Using the slope parameter that was already computed on the surface, an approximation of the B1+ correction can be applied back in the volume, and so we also generate corrected T1w/T2w ratio volumes for all datasets. We note that this approximate correction produces a similar desirable effect to the cortical correction, including reducing the center to peripheral bias and improving left-right symmetry of the white matter (though it does not account for any contrast differences between grey and white matter explicitly). It is not possible to run either the Eq. 6 L-R or Eq. 7 I-T cost functions directly on the volume data because of misalignments of the cortical folds in the volume. For the same reasons, we stress that it is NOT appropriate to attempt across-subject “VBM-style” analyses of these T1w/T2w ratio volumes in lieu of proper surface-based cortical analyses for reasons described elsewhere (Glasser et al., 2016b; Coalson et al., 2018), and also because partial volume effects of CSF, grey matter, and white matter will completely dominate the differential effects of myelin content in traditional volume-based analyses (i.e., the differences in T1w/T2w ratio between tissues are much larger than those within tissues; see Discussion).

## 3. Results

### 3.1. HCP-YA: Group Average T1w/T2w Myelin Map Transmit Field Correction

We begin by illustrating the group average T1w/T2w myelin map (top two rows left) in Figure 5 together with the group average AFI map (top two rows right)^14^. As in Figure 4 the group average AFI map lacks obvious artifacts and can directly be used to correct the group average T1w/T2w myelin map. Importantly, the left-right asymmetry (computed using the asymmetry index in Equation #6) in the T1w/T2w myelin map (bottom row left) and AFI map (bottom row right) are highly correlated spatially (r=0.94) even though different in magnitude (necessitating the fitting procedure described in methods sections 2.1 and 2.7).

**Figure 5.**
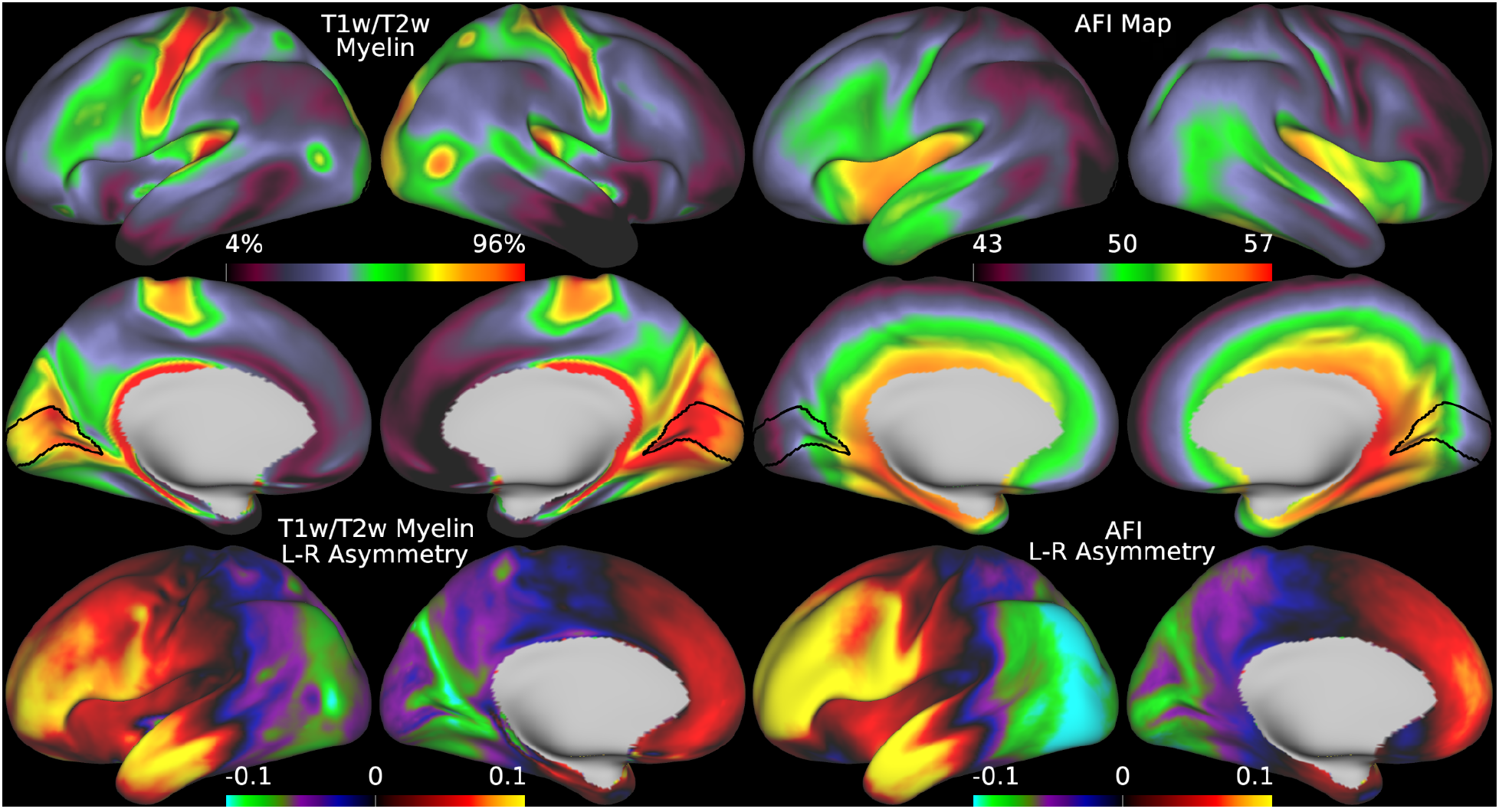
illustrates the group average T1w/T2w myelin maps (top two rows left), group average AFI maps showing the B1+ field as flip angles (top two rows right, in degrees), left-right asymmetry in T1w/T2w myelin maps (bottom row left), and left-right asymmetry in AFI maps (bottom row right). Area V1 is outlined in black. Methods sections 2.2 and 2.3 describe the participants and data, and preprocessing is described in methods sections 2.6 and 2.7. https://balsa.wustl.edu/7qwq3.

Thus, minimizing left-right asymmetry in the group average T1w/T2w myelin map (Equation #6) is an appropriate cost function for optimizing the slope parameter in Equation #5. Figure 6 shows the original and corrected T1w/T2w myelin maps and their asymmetry maps. The correction has markedly reduced the spatial correlation of the left-right asymmetry between the AFI and T1w/T2w myelin maps (r=0.10). Additionally, beyond just correcting hemispherically asymmetric biases, the correction has also addressed *symmetric* biases in the myelin maps. For example, the homogeneity of visual area V1, which contains a strong central to peripheral flip angle gradient (see in Figure 5), is markedly improved by this correction as seen in Figure 6. Correction of both the asymmetric and symmetric biases make the T1w/T2w myelin maps appear more similar to alternative non-invasive myelin measures that have undergone B1+ bias correction, such as quantitative T1, T2*, and MT, which also lack both these asymmetric and symmetric biases (Carey et al., 2018). Finally, there are some real left-right hemispheric differences that are not removed and that are due to slightly different positions of the cortical areas relative to cortical folds after the folding-based registration used to align the FS_LR template across hemispheres (Van Essen et al., 2012), for example the positive/negative couplets near the MT+ complex, LIPv in the intraparietal sulcus, and along the precentral gyrus (see faint circles). Such residual asymmetries show that the transmit field correction is not removing all hemispheric asymmetries from the data.

**Figure 6.**
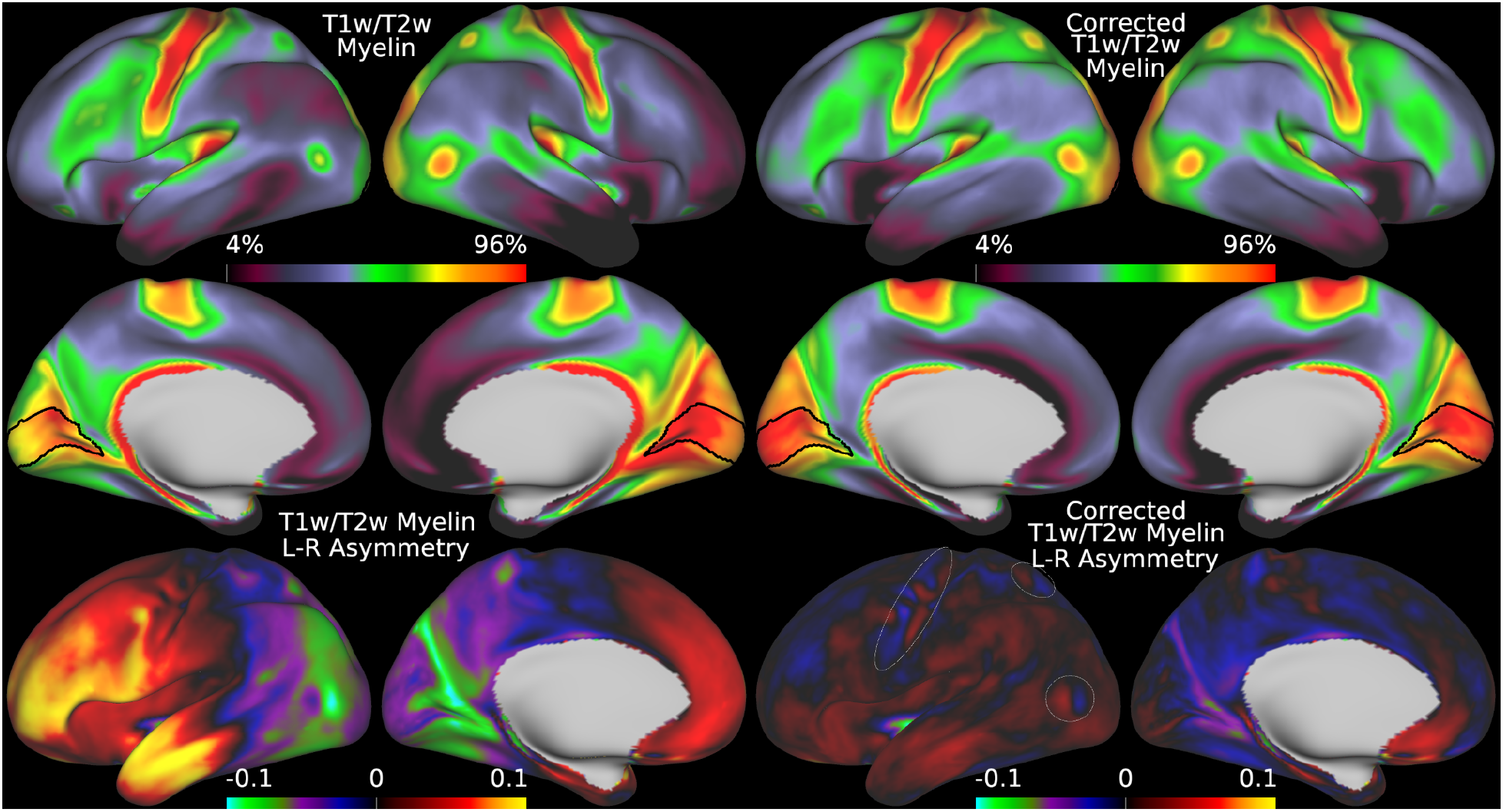
illustrates the group average original T1w/T2w myelin maps (top two rows left), group average corrected T1w/T2w myelin maps (top two rows right), left-right asymmetry in original T1w/T2w myelin maps (bottom row left), and left-right asymmetry in corrected T1w/T2w myelin maps (bottom row right). Area V1 is outlined in black. The faint circles represent the MT+ complex, LIPv, and M1. The L-R cost function (Eq. 6) was used for this correction. Methods sections 2.2 and 2.3 describe the participants and data, and preprocessing is described in methods sections 2.6 and 2.7. https://balsa.wustl.edu/6VjVk.

### 3.2. HCP-YA: Group Average Effects of Transmit Field Correction of Individual T1w/T2w Myelin Maps

Figure 7 shows a comparison of the group corrected myelin map (with the Left-Right cost function; Equation #6) and the mean individual corrected myelin map (with the Individual-Template cost function; Equation #7). These maps are nearly identical (spatial correlation of r=0.99), illustrating that the individual participant correction using Equation #7 instead of Equation #6 with regularized AFI maps is in the aggregate well behaved. The main differences are due to the additional spatial smoothing that occurs on the AFI data in the individual corrections, and the similarity between these two maps shows that left-right asymmetries are not necessary for B1+ transmit field correction once a group reference template has been generated. The effects on the individuals themselves will be illustrated using the toy statistical model.

**Figure 7.**
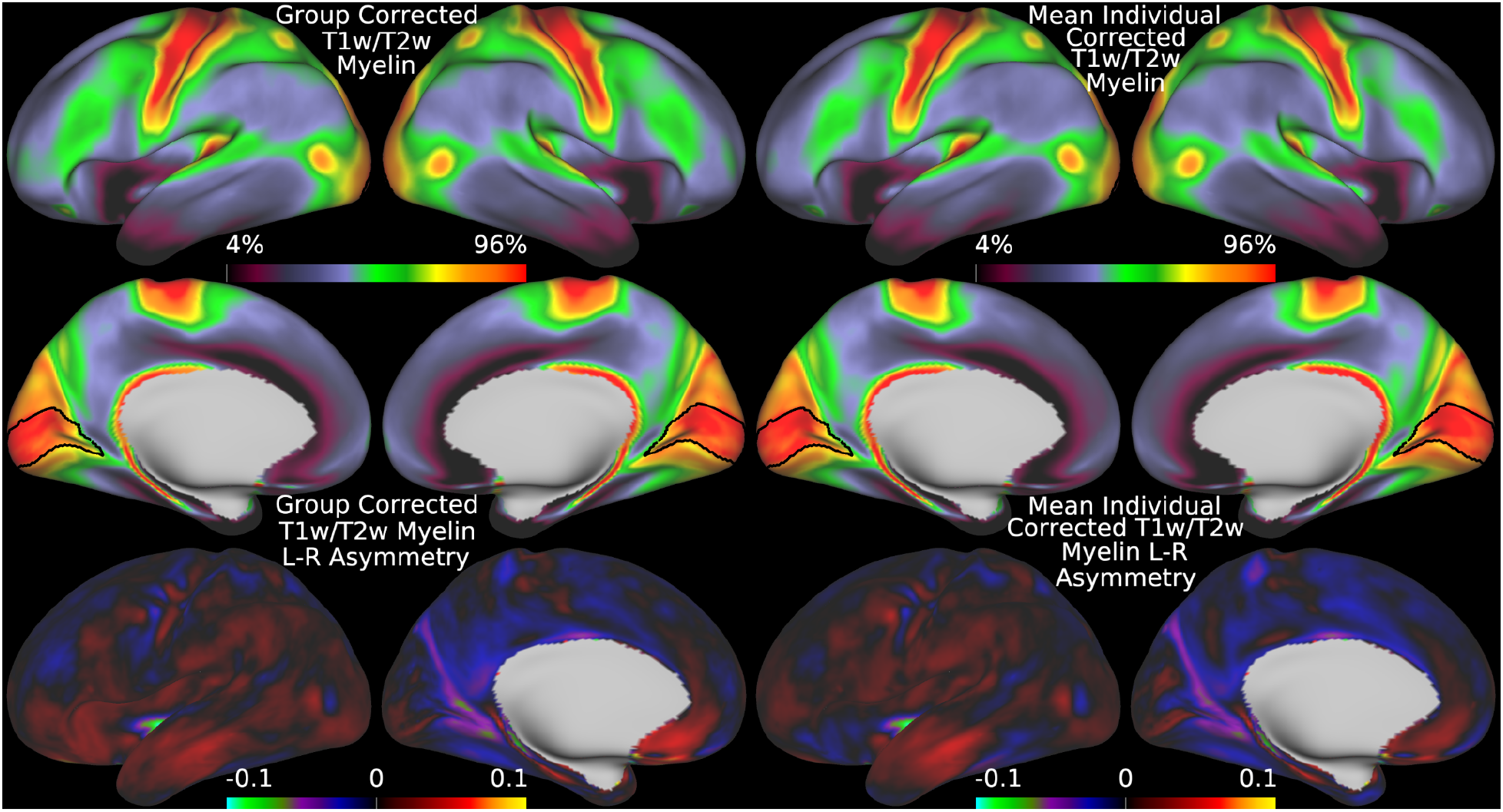
illustrates a comparison between the correction applied to the group average and the mean of the individually corrected myelin maps, showing that they are very similar. The data on the left were corrected with the L-R cost function (Eq. 6) and those on the right were corrected with the I-T cost function (Eq. 7). Methods sections 2.2 and 2.3 describe the participants and data and preprocessing is described in methods sections 2.6 and 2.7. https://balsa.wustl.edu/1BgBP.

### 3.3. HCP-YA: Statistical Effects of Individual T1w/T2w Myelin Map Transmit Field Correction

Although obtaining more accurate group average T1w/T2w myelin maps is useful neuroanatomically, the core goal of this paper is enabling cross-participant analyses that relate T1w/T2w myelin maps to other parameters of interest such as participant demographics, behavior, or disease state. Figure 8 illustrates a multiple linear regression of sex, age, and BMI with the AFI maps, the original T1w/T2w myelin maps, and the B1+ corrected T1w/T2w myelin maps (with the participant-wise covariates of no interest included in the corrected T1w/T2w myelin map regression). This figure demonstrates the sizable effects of subject variables on the transmit field maps, which cause biases in the original T1w/T2w myelin maps, and their elimination in the corrected maps. For example, there is a strong relationship between sex and transmit field that is also present in the original uncorrected maps, suggesting higher T1w/T2w values in men than women across most cortical areas, particularly in the right medial and lateral parietal cortex; however, this effect is eliminated in the corrected T1w/T2w maps where the overall sex differences are quite modest (left 3 columns). This spurious confounding/biasing effect is most likely due to the differences in body and head size, shape, and composition between men and women leading to different flip angles achieved (see Table 1 in the methods), which increases the T1w/T2w ratio in men. Age (middle 3 columns) shows a different set of relationships. In young adults, head size and BMI are only weakly correlated with age (see Table 1). As a result, there is not a strong relationship between age and transmit field (or voltage) across participants (center column in Figure 8). Uncorrected T1w/T2w values show relatively anatomically non-specific increases with age, and correction of the transmit field effects increase this relationship somewhat by reducing nuisance effects of head size and weight/BMI on T1w/T2w myelin maps through the transmit field. Finally, BMI (right 3 columns) has a direct effect on the transmit field due to coil loading. In this figure, BMI shows a strong relationship with both the AFI transmit field and the original T1w/T2w ratio, with a left vs right asymmetry very similar to that illustrated in Figure 5. This effect is dramatically reduced in the corrected T1w/T2w myelin maps.

**Figure 8.**
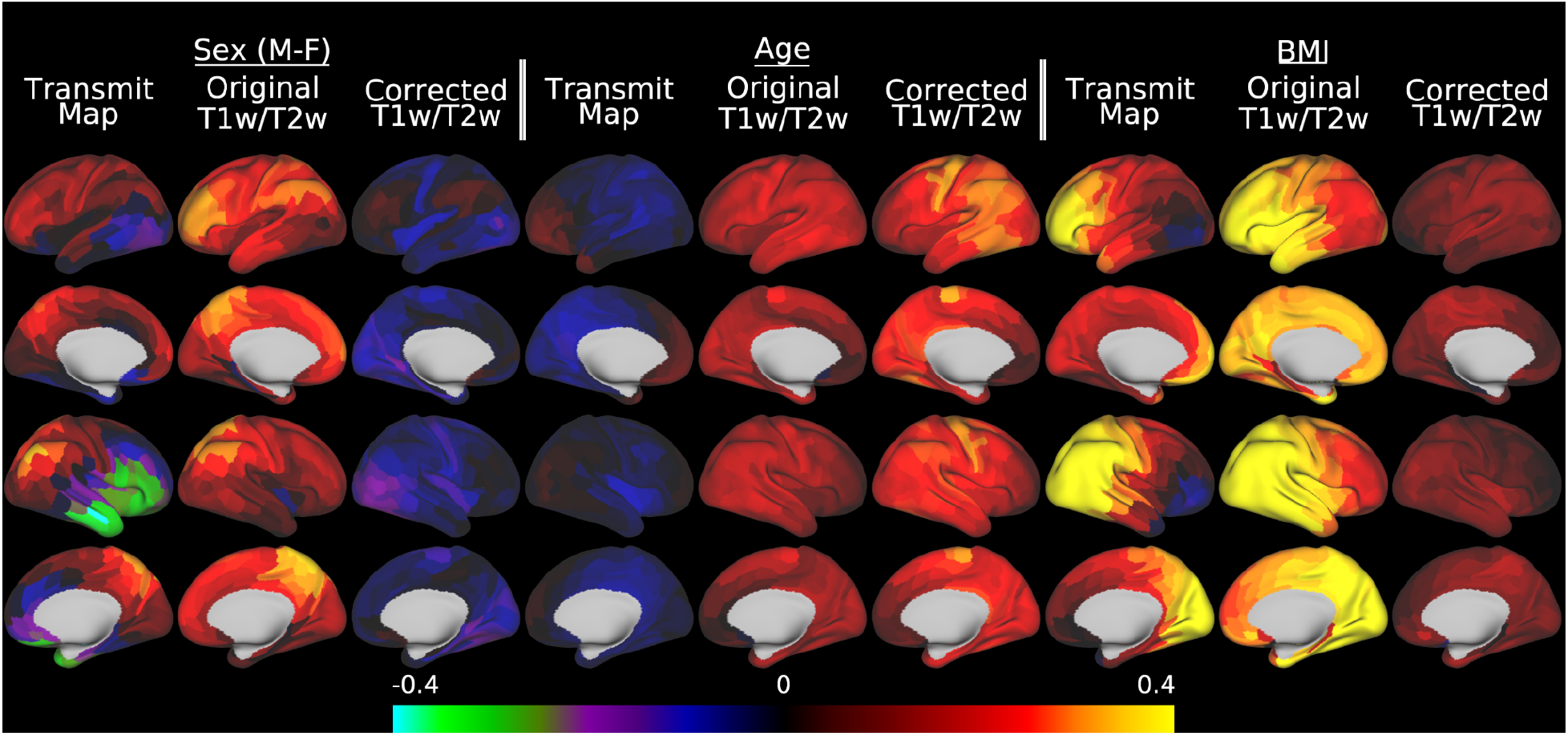
illustrates the effect of transmit field correction of individual T1w/T2w myelin maps on a toy statistical model that includes sex, age, and BMI in a multiple linear regression versus the transmit field maps, the original T1w/T2w myelin maps, and the corrected T1w/T2w myelin maps. Within each grouping by variable of interest, standardized beta values enable comparisons across the AFI transmit maps, original T1w/T2w myelin maps, and the B1+ transmit field corrected T1w/T2w myelin maps. For the corrected maps, the covariates of no interest (including the reference voltage, mean transmit field value, 3 transmit field regularization parameters, and corrected CSF regressor, see Methods section 2.10) are included in the model but not illustrated. All of the data were corrected with the I-T cost function (Eq. 7). Methods sections 2.2 and 2.3 describe the participants and data, preprocessing is described in methods sections 2.6 and 2.7, and the analyses are described in methods sections 2.10 and 2.11. https://balsa.wustl.edu/5XqXP.

### 3.4. HCD: Group Average Effects of Pseudo-Transmit Field Correction of Individual T1w/T2w Myelin Maps

Although we have thus far shown that the transmit field correction approach works when an explicit transmit field has been acquired, many HCP-Style studies have not or are unable to acquire a B1+ map. Thus, we now illustrate the alternative pseudo-transmit field correction approach that can be used in such cases. Figure 9 illustrates that the mean pseudo-transmit field has a similar pattern to the empirically measured (via AFI scans) transmit field in the HCP-YA data, particularly the center versus peripheral bias, and that the HCD data also have left-right asymmetry, though not as pronounced as the HCP-YA data (or as will be shown in the HCA data). Figure 10 demonstrates that individual participant pseudo-transmit field correction using the reference T1w/T2w myelin map generated from the HCP-YA data also successfully removes these asymmetries in HCD data, as in the real transmit field correction in HCP-YA data, and the symmetric biases, producing a more homogeneous area V1.

**Figure 9.**
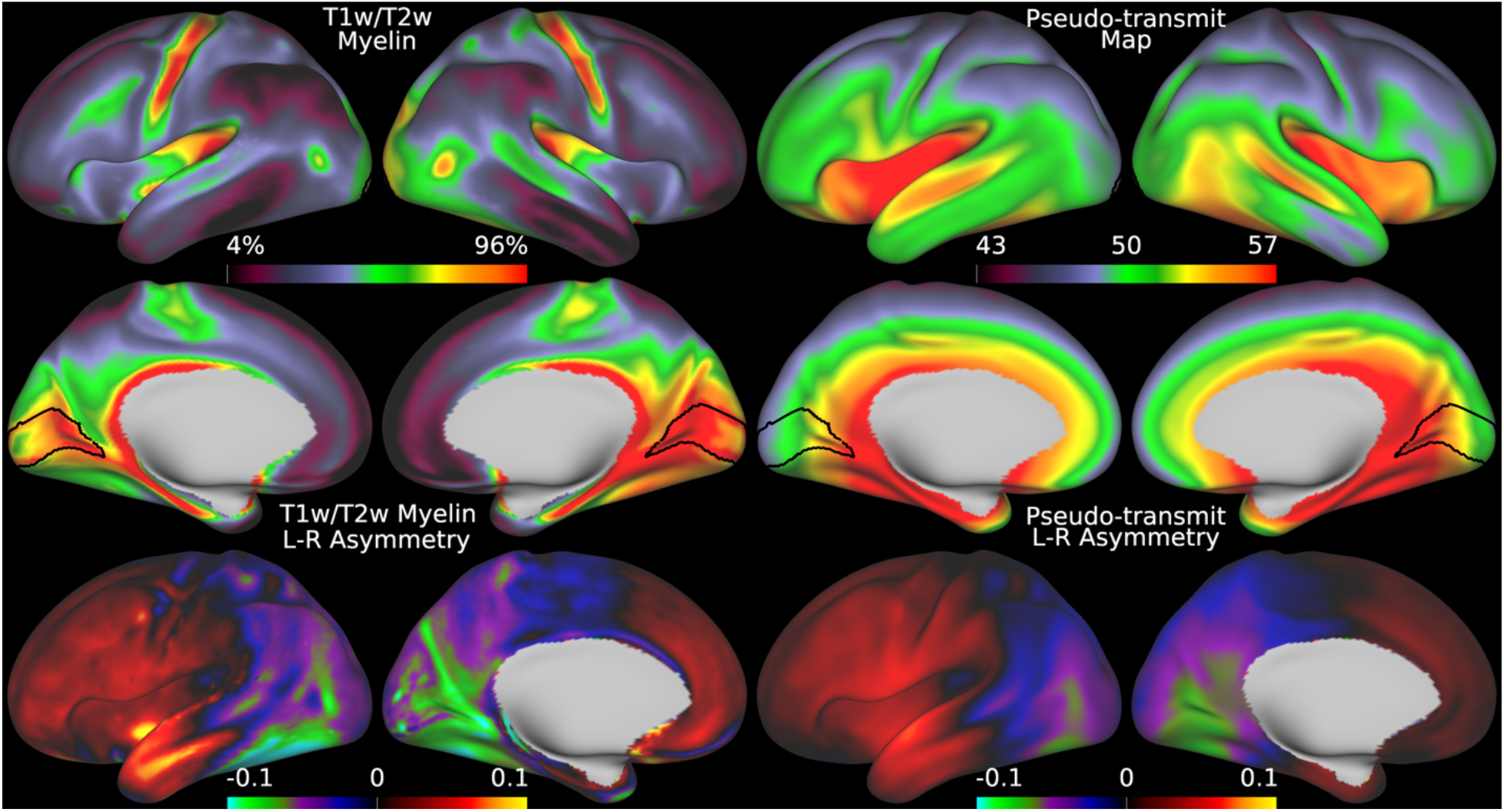
shows the original group average T1w/T2w myelin map from the HCD dataset in the top left two rows together with the regularized group average pseudo-transmit field map in the top right two rows. The units of the pseudo-transmit field map are arbitrary, but are scaled similarly to the transmit field map in the prior figure (5) to ease comparison. The bottom row shows the corresponding asymmetry maps. Methods sections 2.2 and 2.4 describe the participants and data, and preprocessing is described in methods sections 2.6 and 2.8. https://balsa.wustl.edu/nplpK.

**Figure 10.**
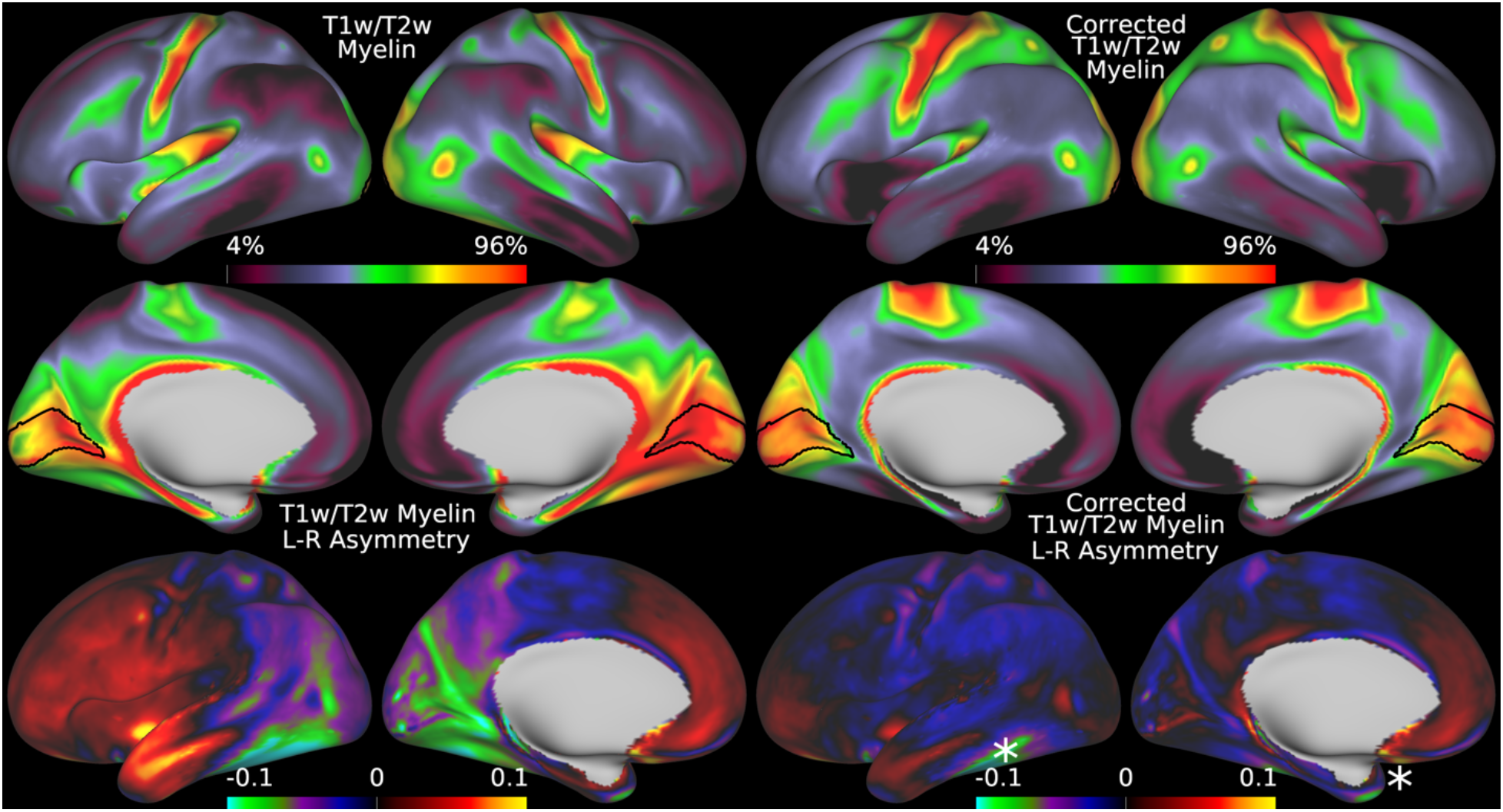
shows the original group average T1w/T2w myelin map from the HCD dataset in the top left two rows together with the mean corrected individual participant T1w/T2w myelin maps. The bottom row shows the corresponding asymmetry maps. The correction works well for most of the brain, but somewhat less well for areas of high susceptibility where the pseudo-transmit field must be imputed from surrounding valid data (marked with stars). The data on the right were corrected with the I-T cost function (Eq. 7). Methods sections 2.2 and 2.4 describe the participants and data, and preprocessing is described in methods sections 2.6 and 2.8. https://balsa.wustl.edu/g767V.

### 3.5. HCD: Statistical Effects of Individual T1w/T2w Myelin Map Pseudo-Transmit Field Correction

Similar to HCP-YA, the toy statistical analysis of sex, age, and BMI contains biases related to head and body size (Figure 11). Similar to HCP-YA, there are regions with differences between boys and girls (e.g., right medial and lateral parietal) in the transmit field map and original uncorrected T1w/T2w myelin maps, which are reduced in the corrected maps, where the sex differences are modest. Although we expect a strong effect of age in developing participants, this is confounded with increasing head and body size (Table 1). The already strong age effect slightly increases after correction, likely through reduction of nuisance effects. Finally, BMI shows the presence of a strong positive but hemispherically asymmetric bias in the transmit field and original maps, as was also the case for HCP-YA. Importantly, BMI in growing adolescents likely represents a different physiological construct than BMI in adults. Regardless of interpretation, however, the salient point here is that the bias is eliminated in the corrected maps.

**Figure 11.**
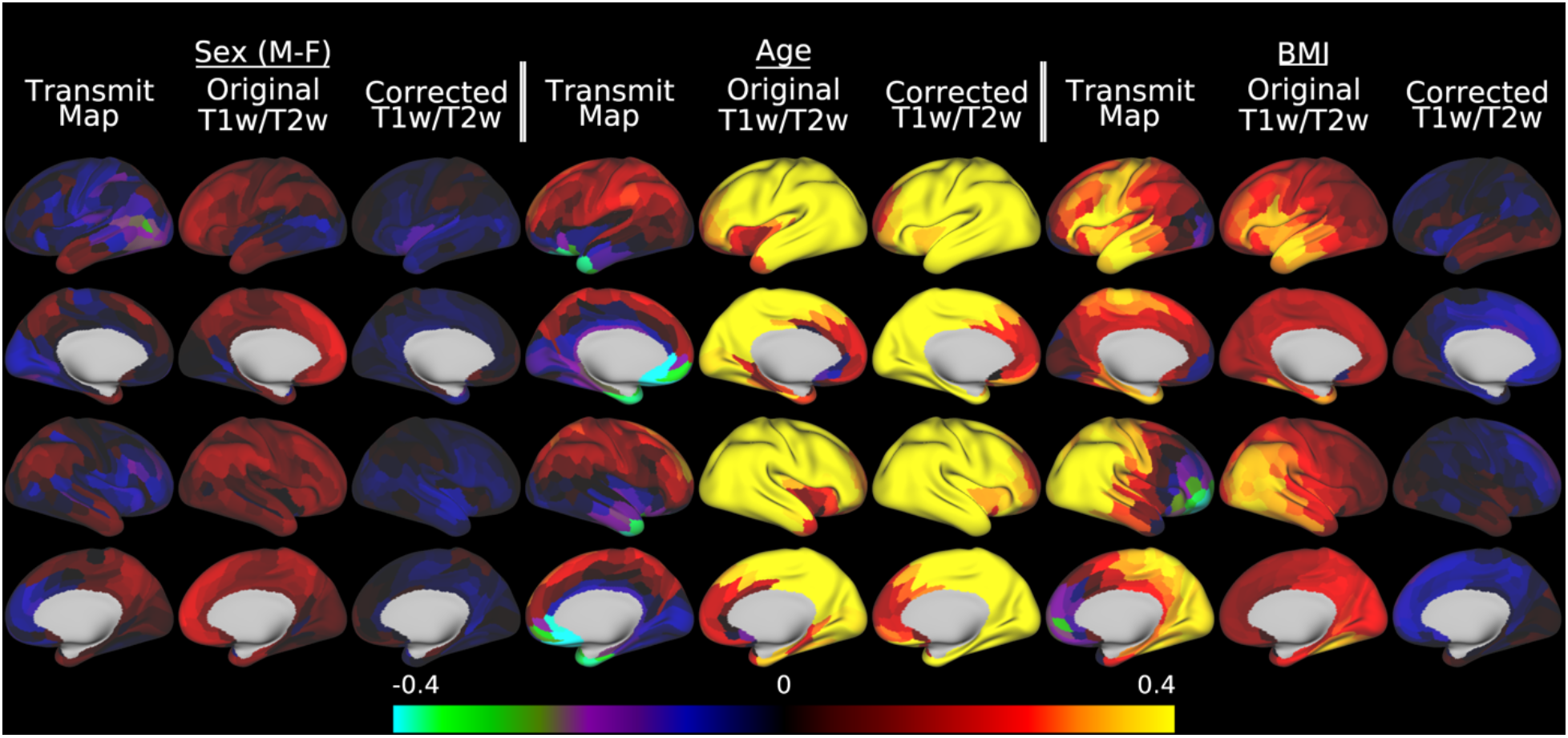
illustrates the effect of transmit field correction of individual T1w/T2w myelin maps on a toy statistical model that includes sex, age, and BMI in a multiple linear regression versus the pseudo-transmit field maps, the original T1w/T2w myelin maps, and the corrected T1w/T2w myelin maps. Within each grouping by variable of interest, standardized beta values enable comparisons across the pseudo-transmit maps, original T1w/T2w myelin maps, and the pseudo-transmit field corrected T1w/T2w myelin maps. For the corrected maps, the covariates of no interest (including the reference voltage, mean transmit field value, 4 transmit field regularization parameters, and corrected CSF regressor, see Methods section 2.10) are included in the model but not illustrated. All of the data were corrected with the I-T cost function (Eq. 7). Methods sections 2.2 and 2.4 describe the participants and data, preprocessing is described in methods sections 2.6 and 2.8, and the analyses are described in methods sections 2.10 and 2.11. https://balsa.wustl.edu/Mxnx8.

### 3.6. HCA: Group Average Effects of Pseudo-Transmit Field Correction of Individual T1w/T2w Myelin Maps

Developing children often have smaller bodies and heads than adults, so a more similar comparator of the pseudo-transmit field approach to the HCP-YA young adults transmit field approach is the HCA aging adults. Figure 12 illustrates that the mean pseudo-transmit field has a similar pattern to the estimated transmit field in the HCP-YA data and the pseudo-transmit field in the HCD data, and that the HCA data also have left-right asymmetry, with the magnitude of the asymmetry lying between the HCP-YA and HCD data, as is expected (MacLennan et al., 2021). Figure 13 demonstrates that individual participant pseudo-transmit field correction using the reference T1w/T2w myelin map generated from the HCP-YA data also successfully removes these asymmetries in the HCA data, as in the HCD data and the real transmit field correction in HCP-YA data, and produces a more homogeneous area V1.

**Figure 12.**
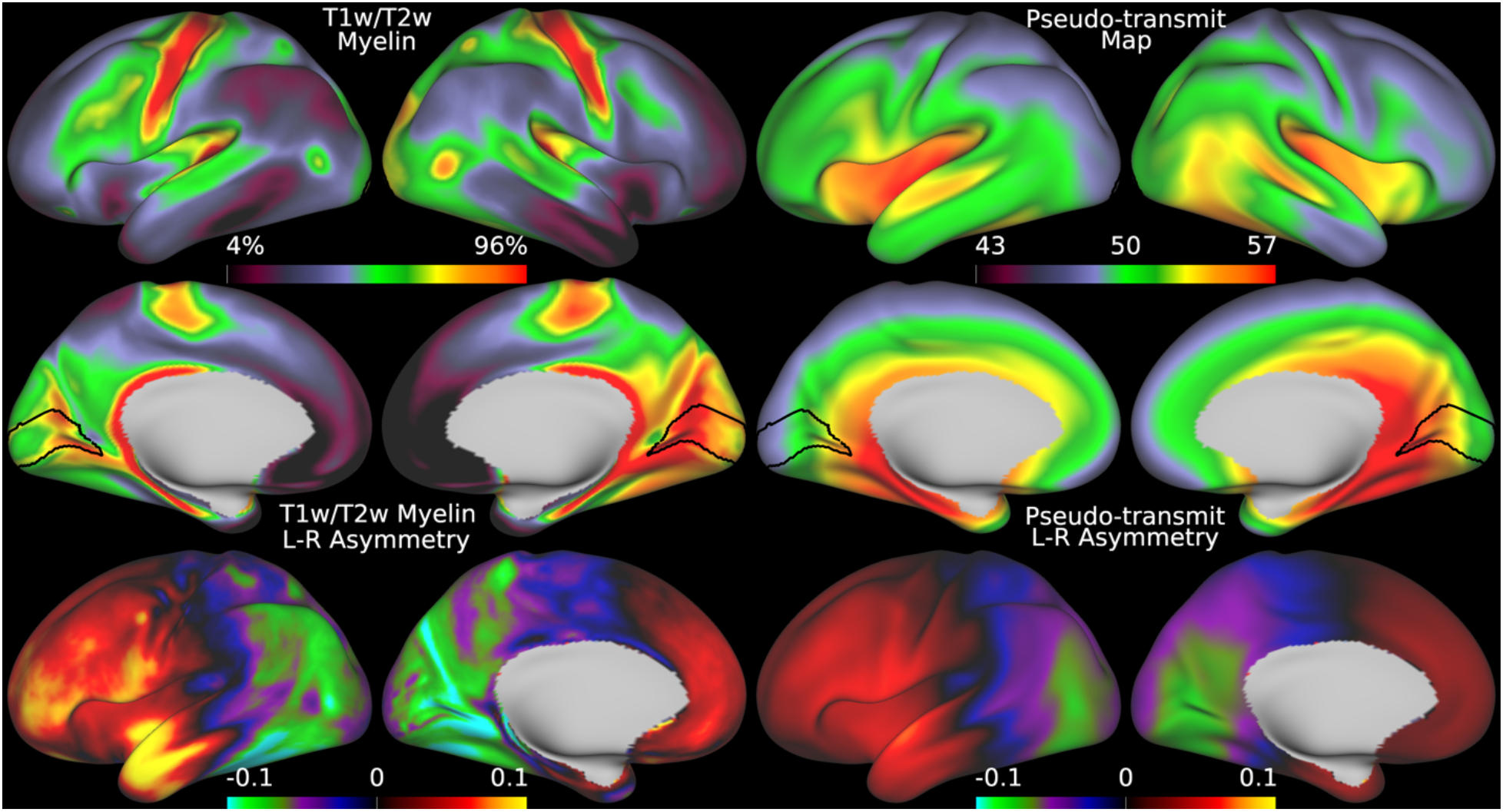
shows the original group average T1w/T2w myelin map from the HCA dataset in the top left two rows together with the regularized group average pseudo-transmit field map in the top right two rows. The units of the pseudo-transmit field map are arbitrary, but are scaled similarly to the transmit field map in the prior figure (5) to ease comparison. The bottom row shows the corresponding asymmetry maps. Methods sections 2.2 and 2.4 describe the participants and data, and preprocessing is described in methods sections 2.6 and 2.8. https://balsa.wustl.edu/B494V.

**Figure 13.**
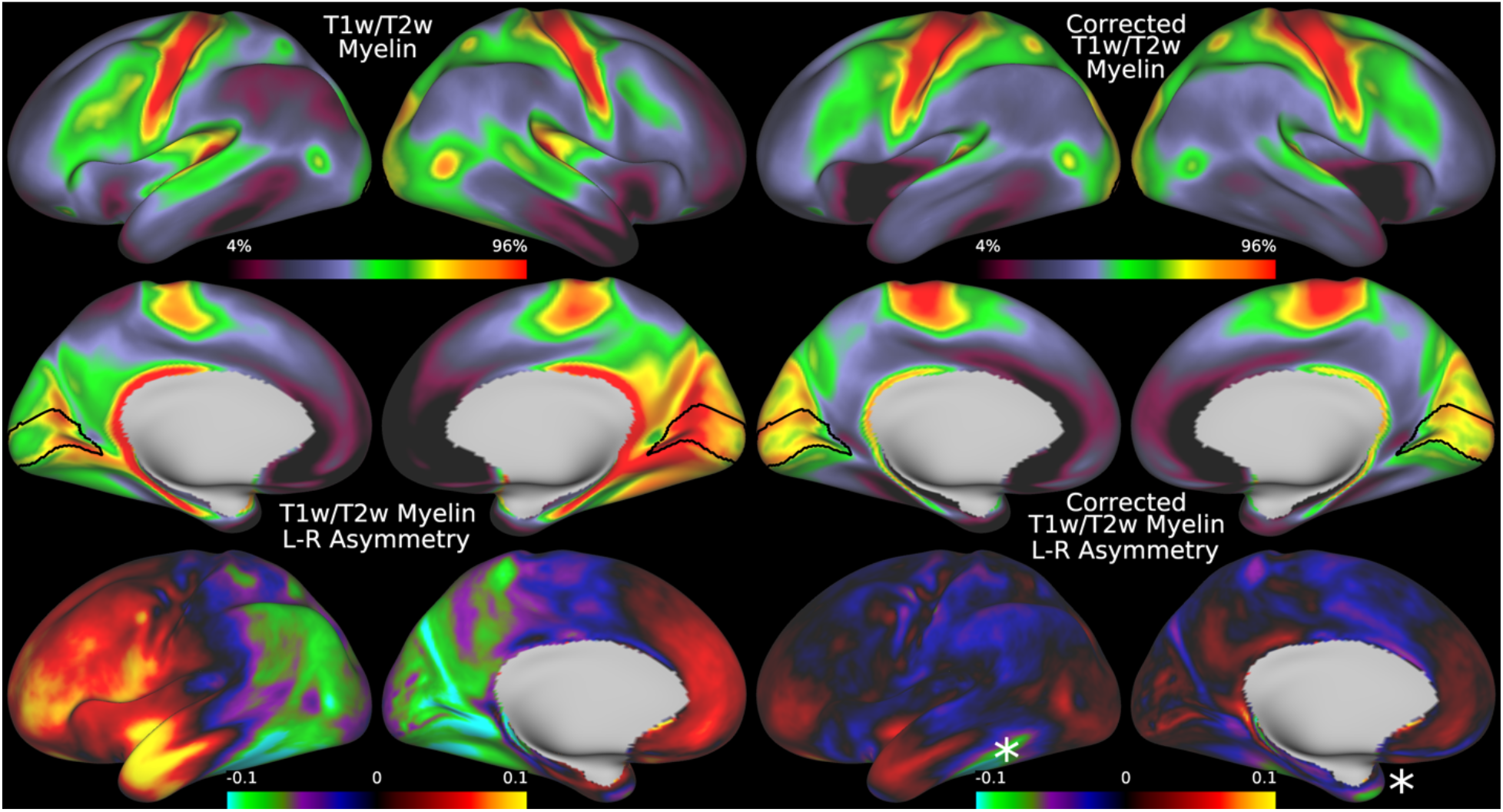
shows the original group average T1w/T2w myelin map from the HCA dataset in the top left two rows together with the mean pseudo-transmit corrected individual participant T1w/T2w myelin maps. The bottom row shows the corresponding asymmetry maps, with stars marking areas of high susceptibility where the pseudo-transmit field must be imputed from surrounding valid data. The data on the right were corrected with the I-T cost function (Eq. 7). Methods sections 2.2 and 2.4 describe the participants and data, and preprocessing is described in methods sections 2.6 and 2.8. https://balsa.wustl.edu/lLMLp.

### 3.7. HCA: Statistical Effects of Individual T1w/T2w Myelin Map Pseudo-Transmit Field Correction

Similar to HCP-YA and HCD, the toy statistical analysis of sex, age, and BMI contains biases related to head and body size (Figure 14). In HCA participants, the correction still improves the symmetry of the maps for sex, age, and BMI (e.g., asymmetry between left and right inferior frontal cortex in the transmit field and original uncorrected T1w/T2w myelin maps between men and women, which is eliminated in the corrected T1w/T2w myelin maps). There is correlation between age and the transmit field map, but again the correction increases the relationship with age, likely by removing nuisance effects. Finally, BMI shows a hemispherically asymmetric pattern in the transmit field and original data that is more similar to HCP-YA than HCD and shows a similar symmetrization in the corrected data, consistent with the strong correlation between BMI and reference voltage (see Table 1 in the methods).

**Figure 14.**
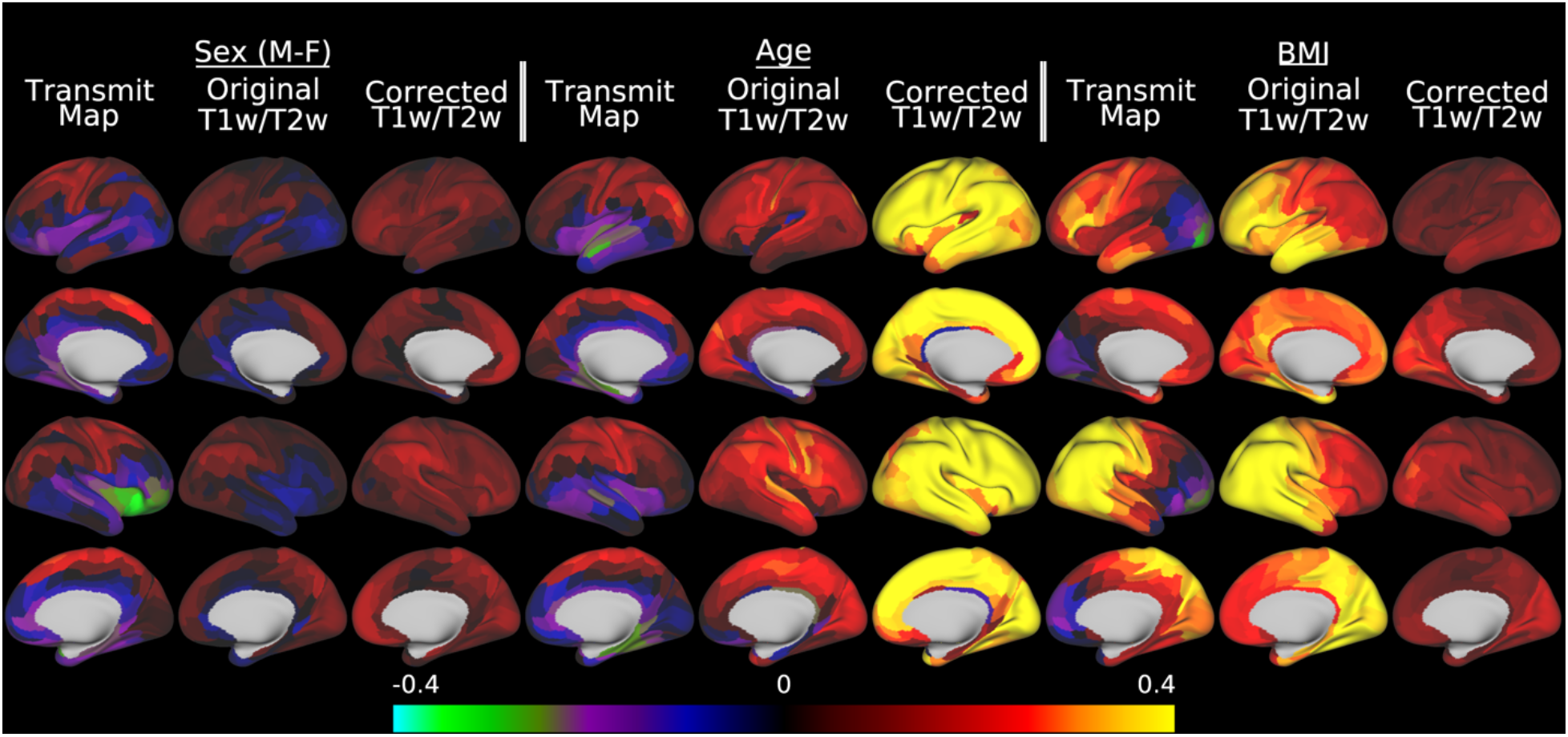
illustrates the effect of transmit field correction of individual T1w/T2w myelin maps on a toy statistical model that includes sex, age, and BMI in a multiple linear regression versus the pseudo-transmit field maps, the original T1w/T2w myelin maps, and the corrected T1w/T2w myelin maps. Within each grouping by variable of interest, standardized beta values enable comparisons across the pseudo-transmit transmit maps, original T1w/T2w myelin maps, and the pseudo-transmit field corrected T1w/T2w myelin maps. For the corrected maps, the covariates of no interest (including the reference voltage, mean transmit field value, 4 transmit field regularization parameters, and corrected CSF regressor, see Methods section 2.10) are included in the model but not illustrated. All of the data were corrected with the I-T cost function (Eq. 7). Methods sections 2.2 and 2.4 describe the participants and data, preprocessing is described in methods sections 2.6 and 2.8, and the analyses are described in methods sections 2.10 and 2.11. https://balsa.wustl.edu/qNwNZ.

### 3.8. NHP_NNP: Group Average T1w/T2w Myelin Map Transmit Field Correction in Macaques

Relative to humans, macaques have much smaller heads and bodies, and as a result the inhomogeneity of flip angles is much lower (Figure 15; flip angle range shown is +3 to −3 degrees in the macaque^15^ versus +7 to −7 degrees in the human in Figure 5). Correspondingly, there is much less hemispheric asymmetry in both the group average T1w/T2w myelin map and the B1+ transmit field map. Although the asymmetry is closer to the noise/artifact floor, making estimation more challenging, it is still present in both images and enables estimation of a species-specific reference template (Figure 16). This map eliminates the left-right asymmetry and a symmetric center > peripheral bias in the macaque myelin map, just as in humans.

**Figure 15.**
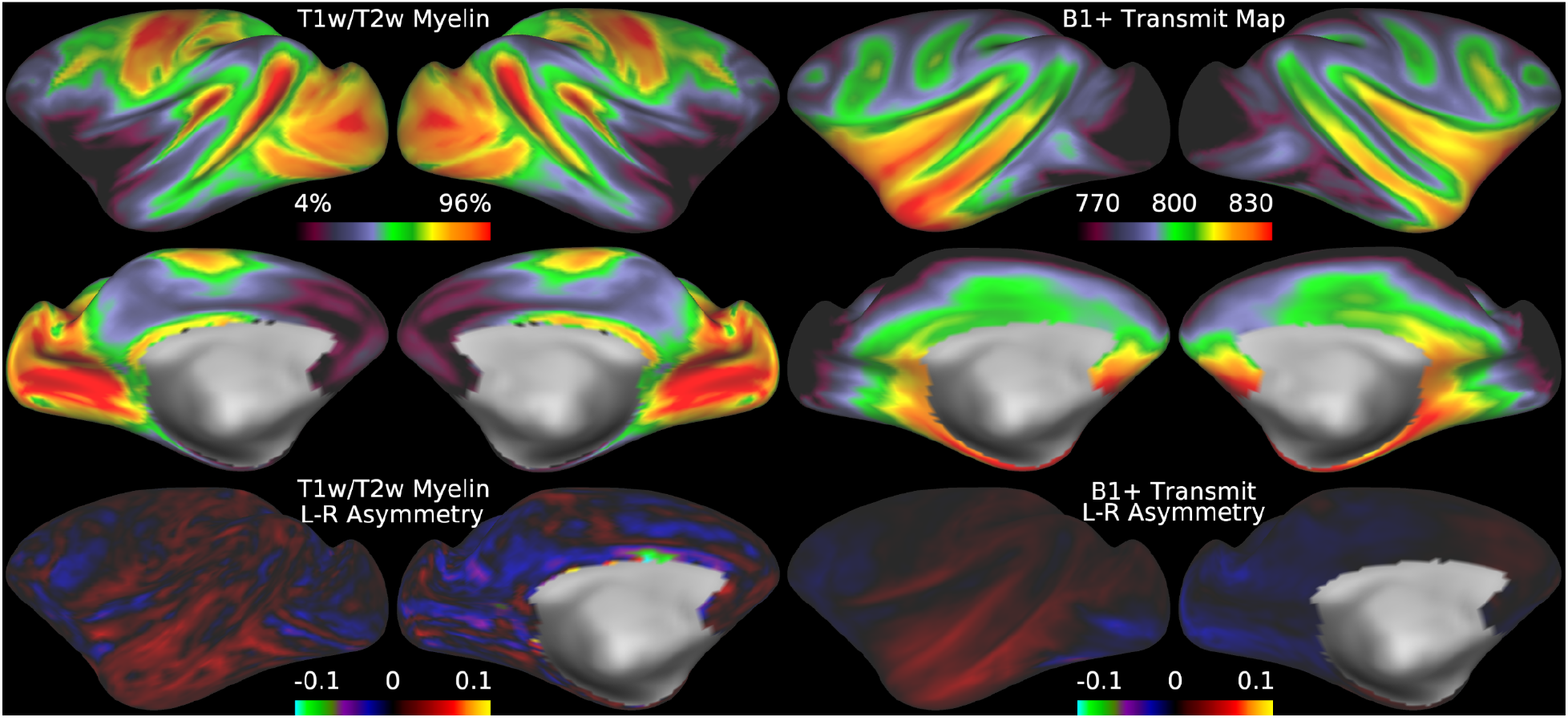
illustrates the group average T1w/T2w myelin maps (top two rows left), group average B1+ transmit field maps (top two rows right), left-right asymmetry in T1w/T2w myelin maps (bottom row left), and left-right asymmetry in group average B1+ transmit field maps (bottom row right). The units of the B1+ transmit field map are flip angle * 10, as generated by the scanner with a reference flip angle of 80 degrees instead of 50 degrees as in humans. Methods sections 2.2 and 2.5 describe the participants and data, and preprocessing is described in methods sections 2.6 and 2.9. https://balsa.wustl.edu/jjnjZ.

**Figure 16.**
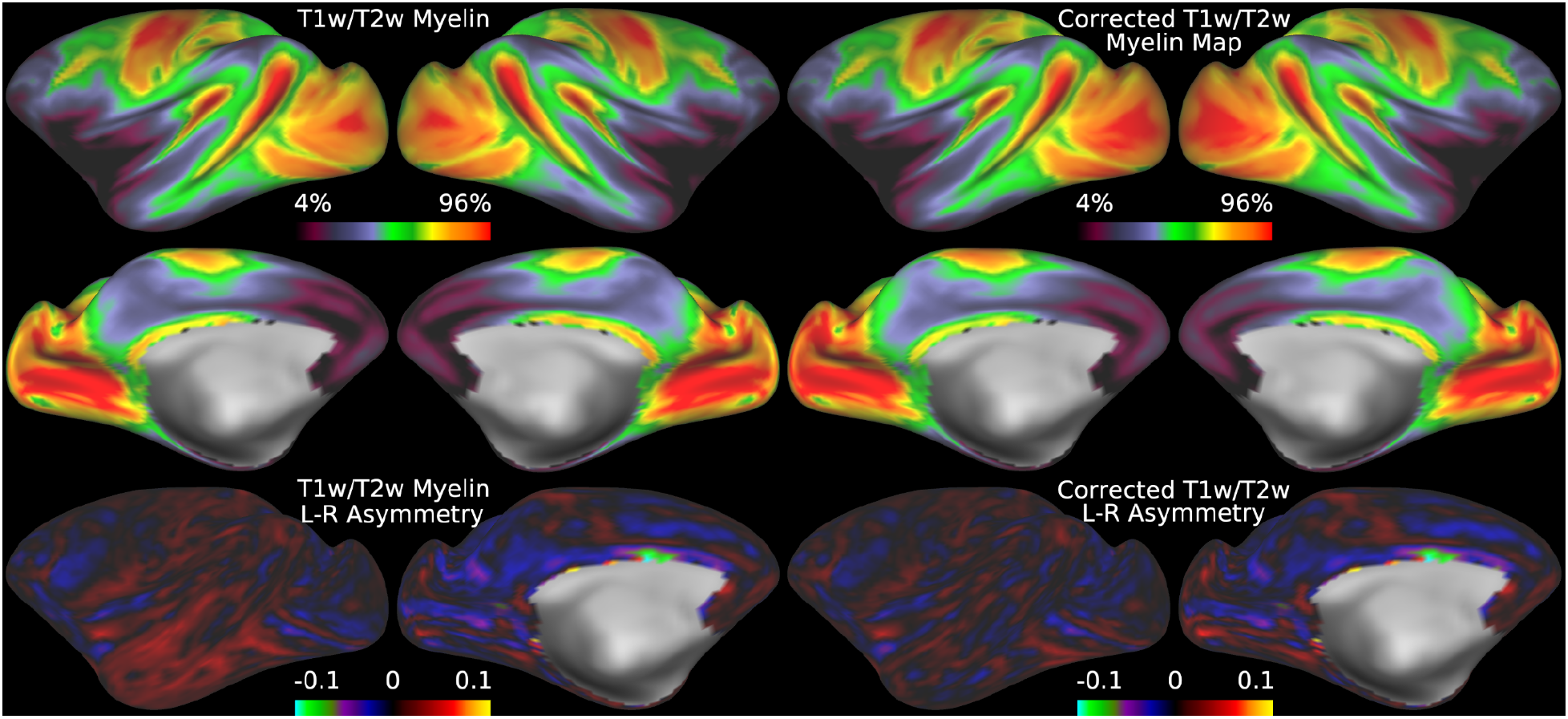
illustrates the group average original T1w/T2w myelin maps (top two rows left), group average B1+ transmit field corrected T1w/T2w myelin maps (top two rows right), left-right asymmetry in original T1w/T2w myelin maps (bottom row left), and left-right asymmetry in corrected T1w/T2w myelin maps (bottom row right). The L-R cost function (Eq. 6) was used for this correction. Methods sections 2.2 and 2.5 describe the participants and data, and preprocessing is described in methods sections 2.6 and 2.9. https://balsa.wustl.edu/wNzNZ.

### 3.9. NHP_NNP: Comparison of Group Average Effects of Transmit and Pseudo-Transmit Field Correction of Individual T1w/T2w Myelin Maps

Macaques are the only dataset in our possession at this time in which both transmit field and pseudo-transmit field data are available for T1w/T2w myelin map correction in the same scanning session, enabling a valuable comparison to be made between them. The spatial correlation between the mean of the individually corrected data and the group reference correction is r=0.999 for the transmit field approach and r=0.995 for the pseudo-transmit field approach. Visually, the mean individual corrected maps are practically indistinguishable (Figure 17) and the mean across participants regularized B1Tx and pseudo-transmit fields correlate at r=0.89.

**Figure 17.**
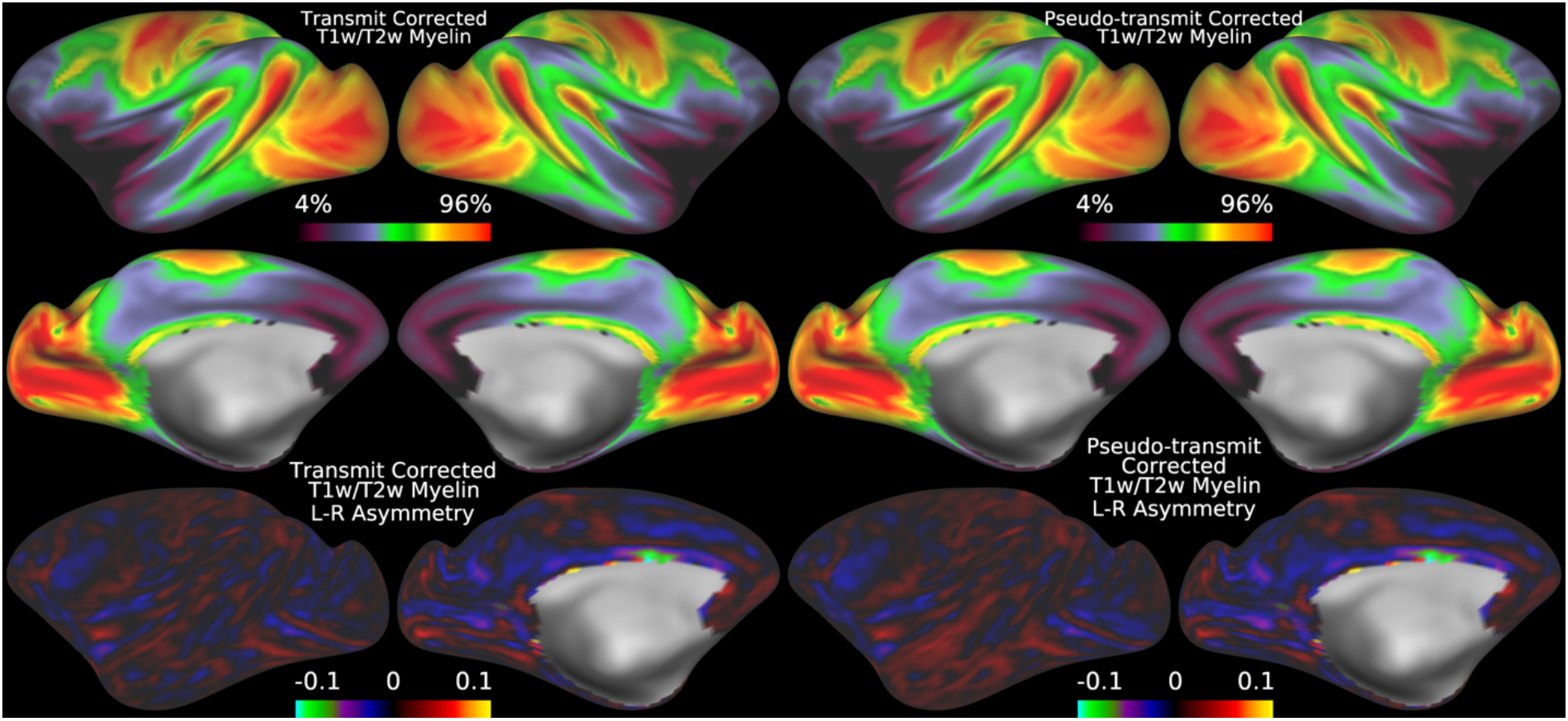
illustrates mean individual transmit field (top left two rows) and pseudo-transmit field (top right two rows) corrected T1w/T2w myelin maps. The corresponding asymmetry maps are shown in the bottom left and right row respectively. All of the data were corrected with the I-T cost function (Eq. 7). Methods sections 2.2 and 2.5 describe the participants and data, and preprocessing is described in methods sections 2.6 and 2.9. https://balsa.wustl.edu/4mlml.

### 3.10. Effects of Transmit and Pseudo-Transmit Field Correction of T1w/T2w Myelin Maps on NHP_NNP Cross-animal Variability and HCP-YA within-participant Reproducibility

The 16-animal macaque dataset is the only one in which we can directly compare the transmit and pseudo-transmit field correction methods for T1w/T2w Myelin Maps. Although the HCP-YA acquired the gradient echo and spin echo EPI data that are prerequisites for the pseudo-transmit correction, it was always in a separate imaging session from the T1w and T2w scans. Thus, it would not be possible to tell whether any differences between transmit and pseudo-transmit field corrections were due to differing effectiveness or simply poorer matching of B1+ fields across different sessions due to differences in head position and/or B1+ homogeneity. Figure 18 illustrates this valuable direct comparison. Eta^2^ is used to quantify similarity (Cohen et al., 2008). Both transmit and pseudo-transmit field corrections improve agreement with the group average data, but transmit field correction appears to perform a bit better (i.e., tighter distribution, higher median). Regressing out the covariates (or including them in the statistical model) further appears to improve both transmit and pseudo-transmit data and makes their performance more similar (medians in the analyses with covariates are closer together, with somewhat tighter distributions in both cases). Thus, we recommend using the covariates (or regressing them out of the maps) depending upon whether one is analyzing a larger statistical model or simply using the maps for downstream neuroanatomical analyses. Although both transmit and pseudo-transmit approaches are reasonable, we recommend explicit transmit field mapping when available scanning time permits its use.

**Figure 18.**
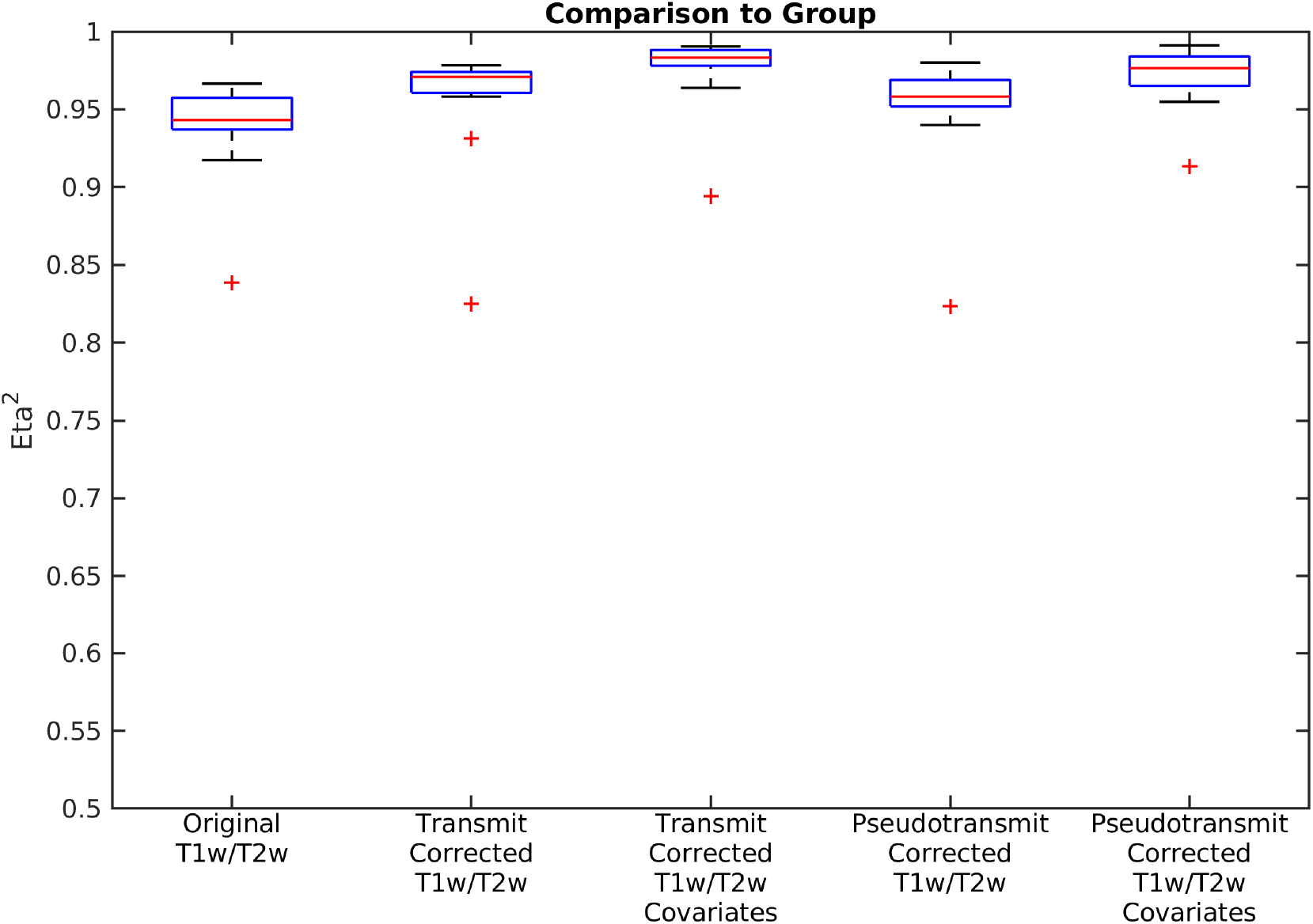
shows boxplots of the eta^2^ between the group average corrected reference T1w/T2w myelin map and the individual T1w/T2w myelin maps for the n=16 macaque datasets. This is shown for the original T1w/T2w myelin values, the transmit field corrected T1w/T2w myelin values without and with covariate regression, and the pseudo-transmit field corrected T1w/T2w myelin values without and with covariate regression. The red line is the median, the edges of the box are the 25th and 75th percentiles (the interquartile range, IQR), the whiskers extend from the end of the IQR to the furthest observation within 1.5 * IQR, and the outliers (+’s) are the data points beyond those limits. The range of eta^2^ is 0-1, with identical inputs yielding 1; inputs that are strictly the negative of each other yielding 0; and values in between represent varying degrees of ‘similarity’ (with 0.5 meaning no relationship). All of the data were corrected with the I-T cost function (Eq. 7). Methods sections 2.2 and 2.5 describe the participants and data, and preprocessing is described in methods sections 2.6, 2.9, 2.10, and 2.12.

The HCP-YA data includes 41 participants in which all modalities used in this study were acquired and processed twice in a test-retest design. Although we would expect the scanner to be fairly reproducible in the B1+ biases it induces in the T1w/T2w individual subjects, the artifacts in the AFI scans and the fitting procedure used to reduce them leave open the possibility that the corrected maps could have worsened reproducibility. Thus, we assess the reproducibility of the data before and after correction with this approach to ensure that reproducibility of T1w/T2w myelin maps is not worsened by transmit field correction^16^. Similar to Figure 18, Figure 19 top row illustrates the improvement in median eta^2^ with transmit field correction and further improvement with covariate regression (or inclusion in the statistical model). A small amount of surface-based smoothing (4mm FWHM) further improves the similarity of the individual data to the group data, as it eliminates random noise and reduces the effects of small surface reconstruction differences. Median eta^2^ is about 0.9 after correction and surface-based smoothing. The bottom row illustrates the test-retest eta^2^ with and without smoothing for each T1w/T2w myelin map and also for unregularized and regularized AFI maps. Reproducibility across visits is similar (medians) without and with the transmit field correction, peaking at median eta^2^ of around 0.9 in smoothed data with correction and covariate regression. The raw AFI maps have markedly reduced reproducibility with median eta^2^ of around 0.76. Regularization improves this median to around 0.9; however. Notably, the transmit coil reference voltages have a test-retest eta^2^ median of 0.95, indicating that the poor test-retest reproducibility of the AFI maps is likely not due to marked differences in scanner B1+ transmit calibration, but rather is chiefly due to the random transmit field artifacts requiring regularization and potentially a smaller component is due to differences in head position inside the body coil (MacLennan et al., 2021). This finding also explains why test-retest reproducibility is similar between corrected and uncorrected data (i.e., the B1+ transmit field effects are reproducible themselves within participants). Overall, it is clear that if AFI is to be used routinely going forward, the artifacts shown here would need to be reduced to produce optimal corrections. If AFI cannot be improved, the B1+ mapping scans used in the macaques would be a better alternative if transmit field correction is desired, given that the B1Tx required less regularization (mean FWHM=13mm) than the AFI (mean FWHM=33mm).

**Figure 19.**
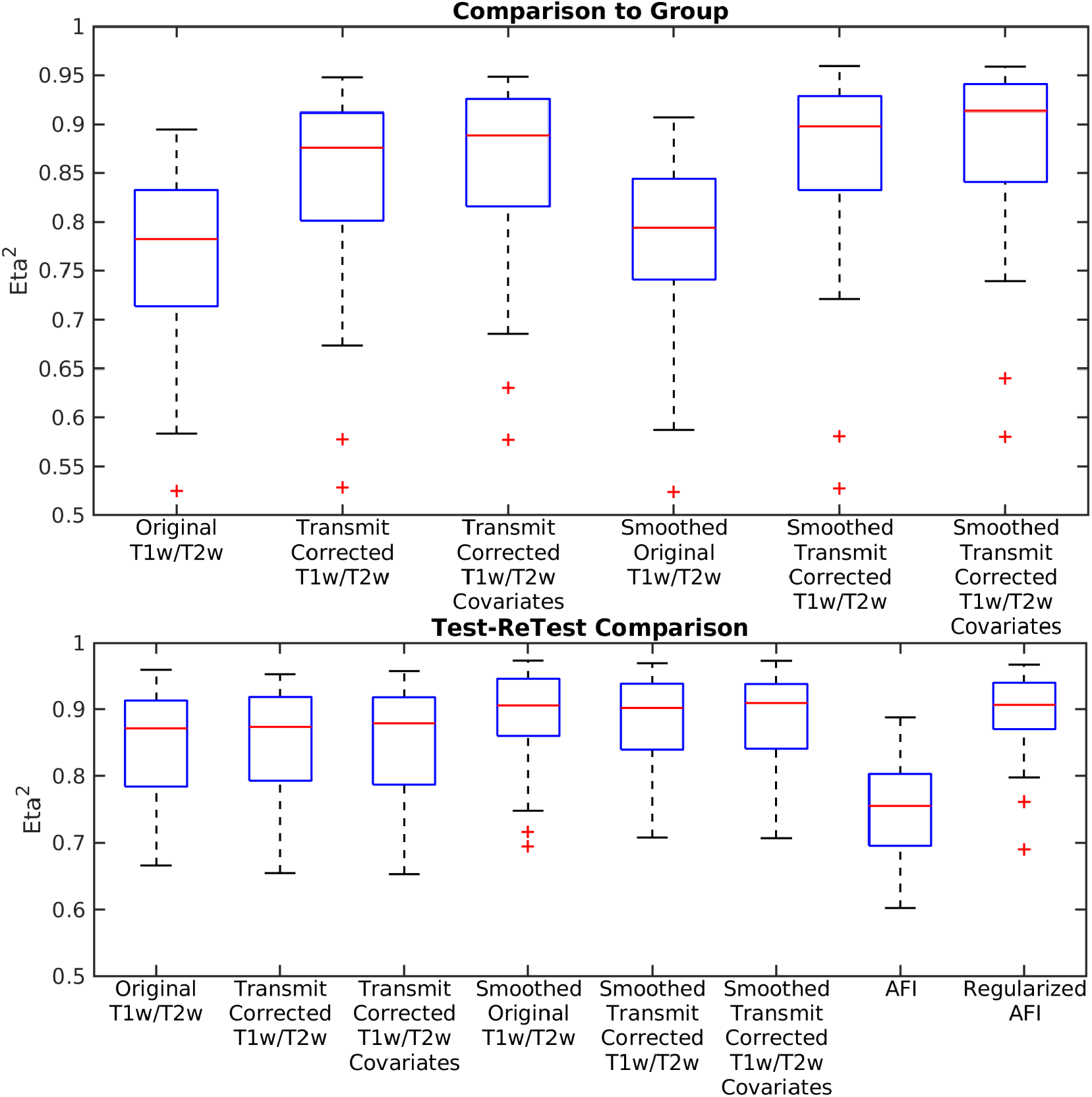
shows box plots across HCP-YA test-retest participants (n=41). Along the top row the eta^2^ between each scan and the reference group average corrected myelin map is illustrated for both the ‘Test’ and ‘ReTest’ visits of the Test-ReTest participants. This is shown for the original T1w/T2w myelin values, the transmit field corrected T1w/T2w myelin values without and with covariate regression, and the transmit field corrected T1w/T2w myelin values without and with covariate regression with smoothing. Along the bottom row, the eta^2^ between the Test and ReTest data is shown for each measure including unregularized and regularized AFI. Smoothing is 4mm FWHM on the surface, which improves test-retest reproducibility somewhat due to reduction in variability from random noise and small surface reconstruction differences. All of the data were corrected with the I-T cost function (Eq. 7). Methods sections 2.2 and 2.3 describe the participants and data, and preprocessing is described in methods sections 2.6, 2.7, 2.10, and 2.12.

## 4. Discussion

We have demonstrated an approximate yet accurate approach to B1+ transmit field correction of T1w/T2w myelin maps using either measured transmit field maps or a derived analog that we term ‘pseudo-transmit’ field maps. We used the finding of highly correlated hemispheric asymmetries between the T1w/T2w cortical myelin maps and B1+ transmit field maps to develop an appropriate correction of both cortical T1w/T2w myelin maps and volume T1w/T2w ratio maps at both the individual and group levels in humans and non-human primates, based on a reasonable set of assumptions (described at the end of the theory section, 2.1, in the methods) that are empirically validated in this study, at least at 3T for the sequences we used. The resulting corrected T1w/T2w myelin maps are largely free of neurobiologically implausible asymmetries, additional symmetric biases, and spurious correlations with head and body size. In addition to being more accurate neuroanatomical representations of the relative differences in myelin content across the cerebral cortex, the transmit field corrected group average maps for each species can be used as reference template maps for future studies’ B1+ corrections of T1w/T2w myelin maps in individuals. Such B1+ corrected T1w/T2w myelin maps are now appropriate for more detailed studies of inter-participant differences with regard to behavior, development, aging, and disease, as well as cross-species differences, some of which will be presented elsewhere (Baum et al., 2021). Importantly, T1w/T2w ratio values are interpretable only as relative differences in a consistently acquired study protocol, and any comparisons across studies would require some kind of harmonization approach. Although this study has focused on Siemens MRI data, these effects are expected to be similar on other vendors’ 3T scanners when a circularly polarized body transmit coil is used for radiofrequency excitation, though empirical validation of these approaches using other vendors’ sequences along the lines presented in this study would be wise.

Interpretation of individual differences in cortical T1w/T2w myelin maps and their relationship to cortical thickness deserves particular attention. The following interpretational guidelines are likely to apply in situations that are not confounded by blurring from motion (which will tend to cause cortex to appear thinner (Reuter et al., 2015) due to errors in surface placement and increased partial voluming of grey matter with CSF and white matter contamination), edema (which will decrease the T1w/T2w ratio), or metal deposition (e.g., iron; which will increase the T1w/T2w ratio)^17^: 1) Apparent increases in cortical T1w/T2w myelin maps if cortical thickness is constant or increasing would likely be due to increases in myelinated axons (e.g., as could occur during development; Bozek et al., 2018). 2) If cortical T1w/T2w measured myelination is increasing at the same time cortical thickness is decreasing, the apparent increase in myelin would likely be due to a decrease in the non-myelinated, plasticity supporting cellular constituents of cerebral cortex (i.e., dendrites, spines, synapses, and glia; Glasser et al., 2014), resulting in a relative increase in myelin density in the remaining cortex (e.g., as may occur during aging), and not necessarily an absolute increase in myelination. 3) Decreases in T1w/T2w with unchanged cortical thickness would likely be due to demyelination (e.g., as could occur in a demyelinating disease). 4) Decreases in T1w/T2w with decreased cortical thickness would likely be due to direct cortical damage (e.g., as in atrophy, such as could occur in a neurodegenerative disease). 5) Decreases in T1w/T2w with increasing thickness are not expected to occur in the absence of pathology (e.g., a neoplasm). Thus, it is helpful to consider what is happening with cortical thickness when interpreting differences in cortical T1w/T2w myelin maps. That said, if image resolution is too low or the surface reconstruction is not optimized, apparent cortical thickness itself can be influenced by cortical myelin content (e.g., Figure 7 auditory core in Glasser and Van Essen 2011) or non-cortical structures such as dura or blood vessels, which can then cause T1w/T2w myelin map artifacts.

Interpretation of volume T1w/T2w maps is more challenging than cortical (surface) T1w/T2w myelin maps, as the relationship between the T1w/T2w MRI contrast and myelin content in the deep white matter has not been shown to be as tight as that seen in cortical grey matter and the true degree of correlation is not yet known (Glasser and Van Essen 2011). Measures of myelin water fraction (MWF) and T1w/T2w ratios are also less correlated in subcortical regions (Arshad et al., 2017 and Uddin et al., 2017), though it is worth noting that MWF measures also show high “myelination” in completely unmyelinated structures such as the dura (Liu et al., 2020). Iron also is less correlated with myelin in subcortical structures than it is in the cerebral cortex (Fukunaga et al., 2010; Stuber et al., 2014; Möller et al., 2019). Figure 20 illustrates the group average B1+ corrected T1w/T2w volume map from the HCP-YA data^18^. Several well-known fiber tracts are visible including the cortico-spinal tract and the optic radiations. Interestingly, while the optic radiations have high T1w/T2w signal, the cortico-spinal tract has relatively low T1w/T2w signal. Also, the motor and visual callosal fibers have relatively lower T1w/T2w signal (a similar pattern is seen with quantitative R1; Harkins et al., 2016). Conversely, the association fibers of the callosum have relatively higher T1w/T2w signal, as does the prefrontal white matter, which is the opposite pattern seen in cortical grey matter (with association areas generally having lower myelin content than primary areas). White matter T1w/T2w signal likely depends on more than the number of myelin wrappings per axon. Additional considerations such as average axonal packing density and average axonal diameter will affect the myelin volume fraction of any given voxel (Stikov et al, 2015). Overall, within the white matter, the T1w/T2w ratio is likely related to the lipid-to-water content ratio, though iron also plays an important role (e.g., see the red nucleus or substantia nigra). That said, as has been seen clinically by neuroradiologists for decades, frank white matter demyelination will result in decreased T1w signal and increased T2w signal leading to a decreased T1w/T2w ratio, and a normally developing young child’s age can be estimated to within a few months based on white matter signal changes found in T1w and T2w images by a skilled pediatric neuroradiologist. Also, as stated previously, cortical myelin content is essentially uninterpretable even in unsmoothed group average T1w/T2w volumes due to cross-participant misalignments of the cerebral cortex, resulting in large partial voluming effects of CSF and white matter that swamp the intracortical grey matter heterogeneities. For this reason, Voxel Based Morphometry (VBM) style analyses are never appropriate in T1w/T2w studies of the cerebral cortex. Skeletonized methods such as Tract-Based-Spatial-Statistics (TBSS; Smith et al., 2006) offer the best option for voxelwise white matter analysis (e.g., Operto et al., 2019), though tractography-based tract analyses would likely be even better (Chen et al., 2017; Thompson et al., 2020).

**Figure 20.**
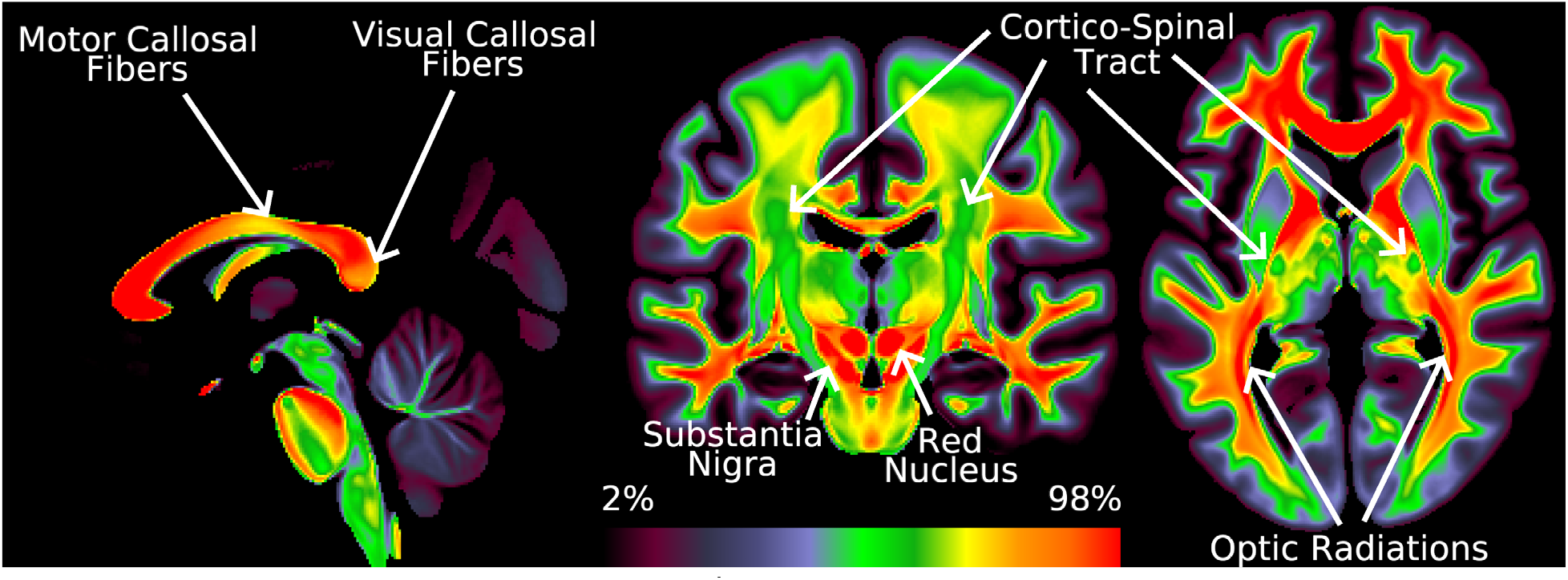
illustrates the group average T1w/T2w map after B1+ correction from the HCP-YA data. The L-R cost function (Eq. 6) was used for this correction. Methods sections 2.2 and 2.3 describe the participants and data, and preprocessing is described in methods sections 2.6., 2.7, and 2.13. https://balsa.wustl.edu/mDBD3. Corresponding uncorrected data is available in the BALSA scene.

There has been debate in the literature as to the optimal approach for non-invasive myelin mapping (Edwards et al., 2018; Campbell et al., 2018; Marques and Norris (2018); Cohen-Adad 2018; Fischl and Sereno 2018; Turner 2019; Shams et al., 2019). In particular, T1w/T2w myelin mapping has been criticized and claimed to not be as specific for myelin as other measures such as quantitative T1, quantitative MT, or T2-based myelin water fraction, or quantitative T2* for iron (Edwards et al., 2018; Arshad et al., 2017; Hagiwara et al., (2018); Uddin et al., 2019). Indeed, T1w and T2w images vary due to a variety of effects within a given voxel, including lipid content, water content, and metal content. If these effects are strong, the T1w/T2w myelin measure will deviate from the actual myelin content in that location of the brain (e.g., vasogenic edema or iron deposition; Yarnykh 2016), though MR tissue contrast properties such as T1, T2*, MT*, or MWF will be similarly affected by these same effects. The T1w/T2w measure has also been criticized relative to other measures such as quantitative T1 owing to difficulties in comparing across protocols and scanners (Edwards et al., 2018), though normalization and harmonization methods have been developed (Ganzetti et al., 2014; Lee et al., 2015). Also importantly, those seeking to directly validate the T1w/T2w measure with histology in the same brains (Righart et al., 2017) have the important challenge that T1w contrast in particular changes early in the post-mortem time period (Fracasso et al., 2016; Shatil et al., 2018; Seifert et al., 2019), making post-mortem T1w/T2w maps a poor surrogate for in vivo T1w/T2w maps (Patel et al., 2020). Despite these criticisms and the fact that quantitative approaches have made some improvements in image resolution and scanning time, T1w/T2w myelin mapping remains the most accessible approach to non-invasive myelin mapping at the time of publication, as it relies on standard product, widely available and routinely used 3D MPRAGE and variable flip angle TSE (e.g., SPACE) sequences on clinical scanners; however, other techniques may become more widely used in the future. Additionally, these high SNR sequences enable high spatial resolution (0.8mm isotropic or better) for precise mapping of the thin cerebral cortex at 3T in clinically reasonable acquisition times (c.f. Trampel et al., 2019; Dong et al., 2021), though future advances may make quantitative approaches as competitive (e.g., Wang et al., 2022). Finally, the T1w and T2w acquisitions are required for making high-quality surface reconstructions used in the HCP-style approach for analyzing fMRI or diffusion images (Glasser et al., 2016b). Thus, we feel that it is desirable to generate and analyze T1w/T2w myelin maps as accurately as possible.

Those choosing to do so should take care to (1) acquire their data with a consistent protocol within a study, (2) acquire their T1w and T2w images at the same spatial resolution (to avoid leaving both resolution and SNR on the table; ideally at 0.8mm isotropic resolution or better) and in close succession (to minimize confounds related to movement between scans), (3) ideally use volumetric navigator 3D sequences (vNav; Tisdall et al., 2012) to minimize the blurring from subject head motion and acquire these scans at the beginning of the session where head motion is often less, (4) acquire a dedicated scan to map the B1+ field if scan time allows (or if not acquire phase reversed gradient echo fMRI and spin echo fieldmaps), and (5) use pre-scan normalize or a vendor equivalent on all images (T1w, T2w, SE, and GRE) to address receive field effects from the head coil in the setting of subject motion.

Investigators who wish to use more quantitative measures might additionally elect to acquire the additional measures for comparison that they consider important such as quantitative T1, quantitative T2*, quantitative magnetization transfer, or T2-based myelin water fraction (e.g., Cohen-Adad et al., 2012; Dick et al., (2012); Sereno et al., 2013; Cohen-Adad (2014); Sánchez-Panchuelo et al., 2014; Mangeat et al., 2015; Liu et al., 2020). Some high-field efforts enable acquiring more than one of these at the same time (Metere et al., 2017; Caan et al., 2019; Sun et al., 2020) or at 3T with longer acquisition times (Carey et al., 2018) and/or lower spatial resolution (Weiskopf et al., 2013). Potential benefits of such quantitative approaches include nominally producing the same values at a given field strength across scanners and sequences without the need for cross-scanner and cross-protocol harmonization and more exact modeling of the MR signal and B1+ correction versus the empirical approximate approaches presented here.

Indeed, the need for B1+ correction of MRI-based myelin maps has long been recognized by proponents of alternative quantitative myelin mapping technologies (e.g., Lutti et al., 2014; Yarnykh 2016; Marques et al., 2017; Hagberg et al., 2017; Haast et al., 2018; Carey et al., 2018). Interestingly, B1+ effects are also visible in certain diffusion measures such as NODDI (Zhang et al., 2012), likely through their effect on the efficiency of spin echo refocusing and thereby SNR (Fukutomi et al., 2018). Of the three approaches used for T1w/T2w myelin map B1+ correction in this study, the B1Tx approach used in the macaque seems to have the fewest artifacts and best overall performance, with a short acquisition time of additional scans (∼ 2 min) and relies on a vendor supplied sequence. The only trick is that even and odd slices must be acquired in separate acquisitions (to fill in the slice gap) and then combined. The pseudo-transmit field approach is likely slightly less accurate due to an increased number of assumptions and methodological complications (as described at the end of the theory section in the methods (2.1), such as the need to threshold out T2* dropout and have similar T2* and T2 contrast) but has the advantage that it allows using images that are already being acquired in a typical HCP-style study. The ringing artifacts in the HCP-YA AFI acquisition could be avoided with adequate spoiling and crusher gradients (Nehrke 2009), which would require careful study/scanner-specific piloting and AFI sequence adjustment. Other approaches for B1+ mapping exist (Desmond et al., 2021), including SA2RAGE (Eggenschwiler et al., 2012), EPI-based approaches (Jiru and Klose 2006), double angle (Cunningham et al., 2006), Bloch-Siegert (Sacolick et al., 2010; Corbin et al., 2019), or improvements on the AFI technique (Maggioni et al., 2021). An optimal B1+ mapping approach should be high resolution (e.g., 2mm isotropic in humans, 1.25mm isotropic in macaques), high SNR, fast to acquire, free of artifacts, and widely available; however, it is out of scope for this study to exhaustively evaluate all of the available methods and make a specific recommendation. Additionally, use of the T1w/T2w ratio at higher field strength (e.g., 7T and above) would require active B1+ shimming with parallel transmit to reduce the range of B1+ inhomogeneity to the more linear range seen at 3T to prevent loss of contrast and/or to satisfy power (i.e., SAR) limitations, particularly in the spin echo T2w image. Further, the longer T1 at higher field strength will increase the required TI and the TR required to achieve a similar TD SNR boosting effect (TD=time to wait before initiating another TR), an effect that can be somewhat offset by the higher SNR at 7T (Bock et al., 2013 Neuroimage), but overall increasing scan time. An alternative high-field approach involves the ratio of two gradient echo images with more similar B1+ effects (i.e., T1w/T2*w; De Martino et al., 2015; T1w/PDw; Van de Moortele et al., (2009)), though T2* has stronger B0 orientation dependence than T2 (Oh et al., 2013) and greater effects from magnetic field inhomogeneity. In summary, we re-emphasize that the success of our empirical correction approach, namely simple multiplication and linear approximation with the transmit field, may be specific to our HCP-style T1w and T2w image acquisition at 3T (see assumption #3 in section 2.1). Generalizing our correction approach to other conditions (e.g., 7T with larger deviations from the investigated acquisition parameters) would benefit from further experiments. Such future extensions could include explicit Bloch equation simulation to map out limits to the parameter space where our assumptions and empirical approximations that work well in this paper are no longer entirely appropriate (e.g., a substantial non-linear contribution) and/or incorporation of non-linear expansions to the correction.

The approaches presented here enable empirical correction of B1+ transmit field effects in T1w/T2w myelin maps at 3T, improving the accuracy of both group and individual maps. These new maps have advantages over the prior “MyelinMap_BC” correction method (Glasser et al., 2013) in that they (1) correct for the *symmetric* biases that were present in the group average and individual maps (e.g., central-to-peripheral bias in intensity) and (2) correct individual-specific biases related to head and body size that may correlate with variables of interest such as age (in development) or BMI, without removing genuine individual differences in T1w/T2w ratio that may be of interest. The corrected ratio maps remain broadly applicable across species (just as the uncorrected or “MyelinMap_BC” corrected maps were), and may in fact enable improved identification of homologous brain regions, as a basis for more precise interspecies cortical surface registration. Prior literature using uncorrected myelin maps or “MyelinMap_BC” corrected myelin maps may need re-evaluation using the new approach depending upon whether results were affected by B1+ effects. Overall, we feel that these correction methods markedly improve the utility of T1w/T2w myelin maps for a wider range of studies when they are appropriate.

## Acknowledgements

Supported in part by HCP-YA: Mapping the Human Connectome: Structure, Function, and Heritability U54MH091657, HCD: Mapping the Human Connectome During Typical Development U01MH109589, HCA: Mapping the Human Connectome During Typical Aging U01AG052564, Connectome Coordination Facility (CCF) I: R24MH108315, Connectome Coordination Facility II: R24MH122820, the McDonnell Center for Systems Neuroscience at Washington University, NIH RO1MH60974 (DCVE, MFG), NIH F30 MH097312 (MFG), NIH T32 EB021955, Brain/MINDS-beyond from Japan Agency of Medical Research and Development (AMED) (JP22dm0307004, P22dm0307006) (TH), and the FreeSurfer Maintenance Grant (R01EB023281) (DG).

## Appendix A

Pseudocode for the pseudo-transmit (PT) field regularization optimization:

~~~
function optim_threshold {
  initialize T2* dropout threshold search range
  while T2* dropout threshold search range is larger than tolerance {
   pick new T2* dropout threshold and apply to PT data
   call optim_smoothing
   update search range
}
  return best result
}

function optim_smoothing {
  initialize smoothing search range
  while smoothing search range is larger than tolerance {
   pick new smoothing amount and apply to PT data
   call optim_slope
  update search range
  }
  return best result
}

function optim_slope {
  initialize slope search range
  while slope search range is larger than tolerance {
   pick new slope value and use it as the correction for myelin (Eq. 5)
   compare corrected myelin to group template (Eq. 7)
   update search range
 }
 return best result
}
~~~

The `while` loop, `pick new value`, and `update search range` operations are all handled via a golden-section search (Kiefer 1953).

## Footnotes

1 The B1+ effects are stronger in HCP-YA data because of the smaller 56cm body transmit coil needed to accommodate the stronger HCP-SC72 gradients (100 mT/m) of the customized Siemens 3T ‘ConnectomS’ platform (Uğurbil et al., 2013). Additionally, these design compromises resulted in the participants’ heads lying 5cm above the magnetic field isocenter and also off-center, relative to the body transmit coil, resulting in reduced B1+ uniformity through the head.

2 Typically, an imaging slab at the center of the scanner.

3 Using a single variable to represent contrast for myelin in both the T1w and T2w images is an approximation, as the contrast properties for the T1w and T2w images related to myelin are not identical (e.g., T1 and T2* in the gradient echo T1w MPRAGE and T1 and T2 in the spin echo T2w SPACE), and of course not all contrast in either image via these MR mechanisms is driven by myelin.

4 The exact equation that Eq. 2 is approximating is substantially more complex, as the spin echo-based T2w SPACE acquisition employs a substantially different RF pulse train (e.g., 90-degree excitation and multiple variable flip angle refocusing pulses up to 180 degrees; Mugler 2014) that propagates differently through the Bloch signal equations than does the gradient echo-based T1w MPRAGE image pulse sequence (e.g., ∼8-degree excitation pulses): 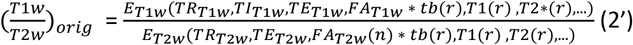 The effect of the transmit bias (tb) field is inherently linked to the tissue contrast and dependent on the MR signal (i.e., Bloch) equations of the underlying acquisition pulse sequences (Mugler and Brookeman 1990; Mugler 2014), where TR, TI, and TE, and flip angle (FA) are specific to the respective acquisition protocol, n indicates multiple variable flip angle refocusing pulses up to 180 degrees in spin echo-based T2w SPACE acquisition, and r represents spatial dependence. The effect of the tb field on T1w/T2w ratio image in its exact form is non-linear (at least quadratic) and related to tissue contrast.

5 As for equation 2, the exact equation that Eq. 3 is approximating is substantially more complex: 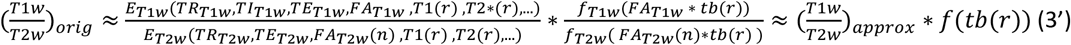 Here we develop an empirical tb field dependent function f(tb(r)), to remove most of the transmit-related intensity bias to yield a bias-corrected T1w/T2w image, (T1w/T2w)_approx_, that approximates the bias-free T1w/T2w image (i.e., (T1w/T2w)_ideal_ below) to a reasonable degree at a desired spatial scale at 3T with these sequences.

6 The T1w scan has the added benefit of using an adiabatic inversion pulse for its initial 180-degree preparation that is designed to be resilient to transmit field inhomogeneities (Garwood and Uğurbil 1992).

7 In the case of the AFI approach in HCP-YA data, the reference flip angle was set to 50 degrees, and to compute TF we divide the flip angle map by 50.

8 The B1+ transmit field is asymmetric in the 3T Siemens Trio, HCP ‘ConnectomS’ (that we will illustrate in Figure 4), and Prisma scanners with circularly polarized body transmit coils. Such an asymmetry exists due to the fundamental dependence of the B1+ field pattern on the electrical properties (i.e., permittivity and conductivity) and geometry (e.g., size) of the sample (Glover et al., 1985; Sled and Pike 1998; Ibrahim et al., 2001), and thus is also expected to be present in other vendors’ scanners as well that use circularly polarized body transmit coils.

9 We chose Golden Search because it is a computationally efficient way to find an optimum of an arbitrary function, and we have used it for this purpose elsewhere in the HCP Pipelines.

10 The gradient echo EPI image experiences an excitation flip angle of ∼50 degrees but the spin echo EPI image experiences an excitation flip angle of 90 degrees and then a 180-degree refocusing pulse.

11 The locations equivalent to the 50-degree spots in the above transmit field maps.

12 Although there are three optimization loops, the computational demands of the pipeline are not high relative to other HCP Pipelines such as FreeSurfer, sICA+FIX, or diffusion preprocessing and all processing for this study was carried out on a small cluster of five 14-core workstations over a few days.

13 The reference flip angle from the Siemens B1+ transmit field phase map is 80 degrees (encoded in the image as flip angle * 10 or 800), so this value was used instead of 50 that was used in the HCP-YA AFI maps in humans.

14 The group myelin map is a simple surface-based average, whereas the TR=20 and TR=120 AFI datasets were separately averaged on the surface before computing the flip angle using the 6th equation from (Yarnykh 2007). This approach is acceptable because the flip angle pattern is generally consistent across participants (same scanner and sequence; MacLennan et al., 2021), the real flip angles form a smooth map, and the nonlinear steps are nearly linear within the range of the data.

15 Units displayed are flip angle * 10 as generated by the scanner with a reference flip angle of 80 degrees instead of 50 degrees as in humans.

16 Note that the purpose of this reproducibility analysis is NOT to neuroanatomically and neurobiologically validate the correction itself, which was done in the preceding sections and figures.

17 Changes across the lifespan or disease states in neocortical MR tissue contrast mechanisms such as T1, T2*, or MT aside from these considerations are presumed to be largely due to changes in myelin density (amount per unit volume); however, the invasive or histological validation of such changes has lagged behind the validation of the larger and more fixed differences in MR tissue contrast across cortical areas (e.g., as in Glasser and Van Essen 2011). That said, gross validation of such changes is provided by comparison of Bozek et al., 2018’s T1w/T2w cortical myelin maps of neonatal development to Flechsig’s classic invasive maps that show the order of myelination in the cerebral cortex during neonatal development (e.g., Von Bonin 1950).

18 B1+ transmit field correction in the volume also improves the symmetry of the white matter T1w/T2w intensities and removes a center versus peripheral bias in T1w/T2w intensities, despite the fact that the fitting procedure is based on the grey matter alone and there theoretically will be an interaction between tissue contrast and B1+ effects, as discussed in Section 2.1.

## References

Arshad, M., Stanley, J.A. and Raz, N., 2017. Test–retest reliability and concurrent validity of in vivo myelin content indices: Myelin water fraction and calibrated T1w/T2w image ratio. Human brain mapping, 38(4), pp.1780–1790.

Autio, J.A., Glasser, M.F., Ose, T., Donahue, C.J., Bastiani, M., Ohno, M., Kawabata, Y., Urushibata, Y., Murata, K., Nishigori, K. and Yamaguchi, M., 2020. Towards HCP-Style macaque connectomes: 24-Channel 3T multi-array coil, MRI sequences and preprocessing. Neuroimage, 215, p.116800.

Baum, G., Flournoy, J., Glasser, M.F, Harms, M., Mair, P., Sanders, A., Barch, D., Buckner, R., Bookheimer, S., Dapretto, M., Smith, S., Thomas, K., Yacoub, E., Van Essen, D., and Somerville, L., 2021. Graded Variation In Cortical T1w/T2w Myelination During Adolescence. BioRXiv.

Bock, N.A., Hashim, E., Janik, R., Konyer, N.B., Weiss, M., Stanisz, G.J., Turner, R. and Geyer, S., 2013. Optimizing T1-weighted imaging of cortical myelin content at 3.0 T. Neuroimage, 65, pp.1–12.

Bonny, J.M., Foucat, L., Laurent, W. and Renou, J.P., 1998. Optimization of Signal Intensity andT1-Dependent Contrast with Nonstandard Flip Angles in Spin-Echo and Inversion-Recovery MR Imaging. Journal of Magnetic Resonance, 130(1), pp.51–57.

Bookheimer, S.Y., Salat, D.H., Terpstra, M., Ances, B.M., Barch, D.M., Buckner, R.L., Burgess, G.C., Curtiss, S.W., Diaz-Santos, M., Elam, J.S. and Fischl, B., 2019. The lifespan human connectome project in aging: an overview. Neuroimage, 185, pp.335–348.

Bozek, J., Makropoulos, A., Schuh, A., Fitzgibbon, S., Wright, R., Glasser, M.F., Coalson, T.S., O’Muircheartaigh, J., Hutter, J., Price, A.N. and Cordero-Grande, L., 2018. Construction of a neonatal cortical surface atlas using Multimodal Surface Matching in the Developing Human Connectome Project. NeuroImage, 179, pp.11–29.

Burt, J.B., Demirtaş, M., Eckner, W.J., Navejar, N.M., Ji, J.L., Martin, W.J., Bernacchia, A., Anticevic, A. and Murray, J.D., 2018. Hierarchy of transcriptomic specialization across human cortex captured by structural neuroimaging topography. Nature neuroscience, 21(9), pp.1251–1259.

Caan, M.W., Bazin, P.L., Marques, J.P., de Hollander, G., Dumoulin, S.O. and van der Zwaag, W., 2019. MP2RAGEME: T1, T2*, and QSM mapping in one sequence at 7 tesla. Human brain mapping, 40(6), pp.1786–1798.

Campbell, J.S., Leppert, I.R., Narayanan, S., Boudreau, M., Duval, T., Cohen-Adad, J., Pike, G.B. and Stikov, N., 2018. Promise and pitfalls of g-ratio estimation with MRI. Neuroimage, 182, pp.80–96.

Carey, D., Caprini, F., Allen, M., Lutti, A., Weiskopf, N., Rees, G., Callaghan, M.F. and Dick, F., 2018. Quantitative MRI provides markers of intra-, inter-regional, and age-related differences in young adult cortical microstructure. Neuroimage, 182, pp.429–440.

Chen, H., Budin, F., Noel, J., Prieto, J.C., Gilmore, J., Rasmussen, J., Wadhwa, P.D., Entringer, S., Buss, C. and Styner, M., 2017, February. White matter Fiber-based analysis of T1w/T2w ratio map. In Medical Imaging 2017: Image Processing (Vol. 10133, p. 101330P). International Society for Optics and Photonics.

Chung, S., Kim, D., Breton, E. and Axel, L., 2010. Rapid B1+ mapping using a preconditioning RF pulse with TurboFLASH readout. Magnetic resonance in medicine, 64(2), pp.439–446.

Collewet, G., Davenel, A., Toussaint, C. and Akoka, S., 2002. Correction of intensity nonuniformity in spin-echo T1-weighted images. Magnetic resonance imaging, 20(4), pp.365–373.

Coalson, T.S., Van Essen, D.C. and Glasser, M.F., 2018. The impact of traditional neuroimaging methods on the spatial localization of cortical areas. Proceedings of the National Academy of Sciences, 115(27), pp.E6356–E6365.

Cohen-Adad, J., 2014. What can we learn from T2* maps of the cortex? Neuroimage, 93, pp.189–200.

Cohen-Adad, J., 2018. Microstructural imaging in the spinal cord and validation strategies. Neuroimage, 182, pp.169–183.

Cohen-Adad, J., Polimeni, J.R., Helmer, K.G., Benner, T., McNab, J.A., Wald, L.L., Rosen, B.R. and Mainero, C., 2012. T2* mapping and B0 orientation-dependence at 7 T reveal cyto-and myeloarchitecture organization of the human cortex. Neuroimage, 60(2), pp.1006–1014.

Cohen, A.L., Fair, D.A., Dosenbach, N.U., Miezin, F.M., Dierker, D., Van Essen, D.C., Schlaggar, B.L. and Petersen, S.E., 2008. Defining functional areas in individual human brains using resting functional connectivity MRI. Neuroimage, 41(1), pp.45–57.

Corbin, N., Acosta-Cabronero, J., Malik, S.J. and Callaghan, M.F., 2019. Robust 3D Bloch-Siegert based mapping using multi-echo general linear modeling. Magnetic resonance in medicine, 82(6), pp.2003–2015.

Cunningham, C.H., Pauly, J.M. and Nayak, K.S., 2006. Saturated double-angle method for rapid B1+ mapping. Magnetic Resonance in Medicine: An Official Journal of the International Society for Magnetic Resonance in Medicine, 55(6), pp.1326–1333.

Delgado, P.R., Kuehne, A., Periquito, J.S., Millward, J.M., Pohlmann, A., Waiczies, S. and Niendorf, T., 2020. B1 inhomogeneity correction of RARE MRI with transceive surface radiofrequency probes. Magnetic Resonance in Medicine, 84(5), pp.2684–2701.

De Martino, F., Moerel, M., Xu, J., van de Moortele, P.F., Ugurbil, K., Goebel, R., Yacoub, E. and Formisano, E., 2015. High-resolution mapping of myeloarchitecture in vivo: localization of auditory areas in the human brain. Cerebral cortex, 25(10), pp.3394–3405.

De Panfilis, C. and Schwarzbauer, C., 2005. Positive or negative blips? The effect of phase encoding scheme on susceptibility-induced signal losses in EPI. Neuroimage, 25(1), pp.112–121.

Desmond, K.L., Xu, R., Sun, Y. and Chavez, S., 2021. A practical method for post-acquisition reduction of bias in fast, whole-brain B1-maps. Magnetic Resonance Imaging, 77, pp.88–98.

Dick, F., Tierney, A.T., Lutti, A., Josephs, O., Sereno, M.I. and Weiskopf, N., 2012. In vivo functional and myeloarchitectonic mapping of human primary auditory areas. Journal of Neuroscience, 32(46), pp.16095–16105.

Deichmann, R., Josephs, O., Hutton, C., Corfield, D.R. and Turner, R., 2002. Compensation of susceptibility-induced BOLD sensitivity losses in echo-planar fMRI imaging. Neuroimage, 15(1), pp.120–135.

Dong, Z., Wang, F., Chan, K.S., Reese, T.G., Bilgic, B., Marques, J.P. and Setsompop, K., 2021. Variable flip angle echo planar time-resolved imaging (vFA-EPTI) for fast high-resolution gradient echo myelin water imaging. NeuroImage, 232, p.117897.

Du, G., Lewis, M.M., Sica, C., Kong, L. and Huang, X., 2019. Magnetic resonance T1w/T2w ratio: A parsimonious marker for Parkinson disease. Annals of neurology, 85(1), pp.96–104.

Edwards, L.J., Kirilina, E., Mohammadi, S. and Weiskopf, N., 2018. Microstructural imaging of human neocortex in vivo. Neuroimage, 182, pp.184–206.

Eggenschwiler, F., Kober, T., Magill, A.W., Gruetter, R. and Marques, J.P., 2012. SA2RAGE: A new sequence for fast B1+-mapping. Magnetic resonance in medicine, 67(6), pp.1609–1619.

Elam, J.S., Glasser, M.F., Harms, M.P., Sotiropoulos, S.N., Andersson, J.L., Burgess, G.C., Curtiss, S.W., Oostenveld, R., Larson-Prior, L.J., Schoffelen, J.M. and Hodge, M.R., 2021. The Human Connectome Project: A retrospective. NeuroImage, p.118543.

Fischl, B., 2012. FreeSurfer. Neuroimage, 62(2), pp.774–781.

Fischl, B. and Sereno, M.I., 2018. Microstructural parcellation of the human brain. Neuroimage, 182, pp.219–231.

Fracasso, A., van Veluw, S.J., Visser, F., Luijten, P.R., Spliet, W., Zwanenburg, J.J., Dumoulin, S.O. and Petridou, N., 2016. Lines of Baillarger in vivo and ex vivo: Myelin contrast across lamina at 7 T MRI and histology. NeuroImage, 133, pp.163–175.

Fukunaga, M., Li, T.Q., van Gelderen, P., de Zwart, J.A., Shmueli, K., Yao, B., Lee, J., Maric, D., Aronova, M.A., Zhang, G. and Leapman, R.D., 2010. Layer-specific variation of iron content in cerebral cortex as a source of MRI contrast. Proceedings of the National Academy of Sciences, 107(8), pp.3834–3839.

Fukutomi, H., Glasser, M.F., Zhang, H., Autio, J.A., Coalson, T.S., Okada, T., Togashi, K., Van Essen, D.C. and Hayashi, T., 2018. Neurite imaging reveals microstructural variations in human cerebral cortical gray matter. Neuroimage, 182, pp.488–499.

Ganzetti, M., Wenderoth, N. and Mantini, D., 2014. Whole brain myelin mapping using T1-and T2-weighted MR imaging data. Frontiers in human neuroscience, 8, p.671.

Gao, R., van den Brink, R.L., Pfeffer, T. and Voytek, B., 2020. Neuronal timescales are functionally dynamic and shaped by cortical microarchitecture. Elife, 9, p.e61277.

Garwood, M. and Uǧurbil, K., 1992. B 1 insensitive adiabatic RF pulses. In In-Vivo Magnetic Resonance Spectroscopy I: Probeheads and Radiofrequency Pulses Spectrum Analysis (pp. 109–147). Springer, Berlin, Heidelberg.

Glasser, M.F., Coalson, T.S., Bijsterbosch, J.D., Harrison, S.J., Harms, M.P., Anticevic, A., Van Essen, D.C. and Smith, S.M., 2018. Using temporal ICA to selectively remove global noise while preserving global signal in functional MRI data. Neuroimage, 181, pp.692–717.

Glasser, M.F., Coalson, T.S., Robinson, E.C., Hacker, C.D., Harwell, J., Yacoub, E., Ugurbil, K., Andersson, J., Beckmann, C.F., Jenkinson, M. and Smith, S.M. and Van Essen, D.C., 2016a. A multi-modal parcellation of human cerebral cortex. Nature, 536(7615), pp.171–178.

Glasser, M.F., Goyal, M.S., Preuss, T.M., Raichle, M.E. and Van Essen, D.C., 2014. Trends and properties of human cerebral cortex: correlations with cortical myelin content. Neuroimage, 93, pp.165–175.

Glasser, M.F., Smith, S.M., Marcus, D.S., Andersson, J.L., Auerbach, E.J., Behrens, T.E., Coalson, T.S., Harms, M.P., Jenkinson, M., Moeller, S. and Robinson, E.C., Sotiropoulos, S.N., Xu, J., Yacoub, E., Ugurbil, K. and Van Essen, D.C., 2016b. The human connectome project’s neuroimaging approach. Nature neuroscience, 19(9), pp.1175–1187.

Glasser, M.F., Sotiropoulos, S.N., Wilson, J.A., Coalson, T.S., Fischl, B., Andersson, J.L., Xu, J., Jbabdi, S., Webster, M., Polimeni, J.R., Van Essen, D.C. and Jenkinson, M., 2013. The minimal preprocessing pipelines for the Human Connectome Project. Neuroimage, 80, pp.105–124.

Glasser, M.F. and Van Essen, D.C., 2011. Mapping human cortical areas in vivo based on myelin content as revealed by T1-and T2-weighted MRI. Journal of Neuroscience, 31(32), pp.11597–11616.

Glover, G.H., Hayes, C.E., Pelc, N.J., Edelstein, W.A., Mueller, O.M., Hart, H.R., Hardy, C.J., O’donnell, M. and Barber, W.D., 1985. Comparison of linear and circular polarization for magnetic resonance imaging. Journal of Magnetic Resonance (1969), 64(2), pp.255–270.

Granberg, T., Fan, Q., Treaba, C.A., Ouellette, R., Herranz, E., Mangeat, G., Louapre, C., Cohen-Adad, J., Klawiter, E.C., Sloane, J.A. and Mainero, C., 2017. In vivo characterization of cortical and white matter neuroaxonal pathology in early multiple sclerosis. Brain, 140(11), pp.2912–2926.

Greve, D.N. and Fischl, B., 2009. Accurate and robust brain image alignment using boundary-based registration. Neuroimage, 48(1), pp.63–72.

Grydeland, H., Vértes, P.E., Váša, F., Romero-Garcia, R., Whitaker, K., Alexander-Bloch, A.F., Bjørnerud, A., Patel, A.X., Sederevičius, D., Tamnes, C.K. and Westlye, L.T., 2019. Waves of maturation and senescence in micro-structural MRI markers of human cortical myelination over the lifespan. Cerebral Cortex, 29(3), pp.1369–1381.

Grydeland, H., Walhovd, K.B., Tamnes, C.K., Westlye, L.T. and Fjell, A.M., 2013. Intracortical myelin links with performance variability across the human lifespan: results from T1-and T2-weighted MRI myelin mapping and diffusion tensor imaging. Journal of Neuroscience, 33(47), pp.18618–18630.

Grydeland, H., Westlye, L.T., Walhovd, K.B. and Fjell, A.M., 2016. Intracortical Posterior Cingulate Myelin Content Relates to Error Processing: Results from T 1-and T 2-Weighted MRI Myelin Mapping and Electrophysiology in Healthy Adults. Cerebral cortex, 26(6), pp.2402–2410.

Haast, R.A., Ivanov, D. and Uludağ, K., 2018. The impact of correction on MP2RAGE cortical T 1 and apparent cortical thickness at 7 T. Human brain mapping, 39(6), pp.2412–2425.

Hagberg, G.E., Bause, J., Ethofer, T., Ehses, P., Dresler, T., Herbert, C., Pohmann, R., Shajan, G., Fallgatter, A., Pavlova, M.A. and Scheffler, K., 2017. Whole brain MP2RAGE-based mapping of the longitudinal relaxation time at 9.4 T. Neuroimage, 144, pp.203–216.

Hagiwara, A., Hori, M., Kamagata, K., Warntjes, M., Matsuyoshi, D., Nakazawa, M., Ueda, R., Andica, C., Koshino, S., Maekawa, T. and Irie, R., 2018. Myelin measurement: Comparison between simultaneous tissue relaxometry, magnetization transfer saturation index, and T 1 w/T 2 w ratio methods. Scientific reports, 8(1), pp.1–12.

Harkins, K.D., Xu, J., Dula, A.N., Li, K., Valentine, W.M., Gochberg, D.F., Gore, J.C. and Does, M.D., 2016. The microstructural correlates of T1 in white matter. Magnetic resonance in medicine, 75(3), pp.1341–1345.

Harms, M.P., Somerville, L.H., Ances, B.M., Andersson, J., Barch, D.M., Bastiani, M., Bookheimer, S.Y., Brown, T.B., Buckner, R.L., Burgess, G.C., Coalson, T.S., Chappell, M.A., Dapretto, M., Douaud, G., Fischl, B., Glasser, M.F., Greve, D.N., Hodge, C., Jamison, K.W., Jbabdi, S., Kandala, S., Li, X., Mair, R.W., Mangia, S., Marcus, D., Mascali, D., Moeller, S., Nichols, T.E., Robinson, E.C., Salat, D.H., Smith, S.M., Sotiropoulos, S.N., Terpstra, M., Thomas, K.M., Tisdall, M.D., Ugurbil, K., van der Kouwe, A., Woods, R.P., Zöllei, L., Van Essen, D.C. and Yacoub, E., 2018. Extending the Human Connectome Project across ages: Imaging protocols for the Lifespan Development and Aging projects. Neuroimage, 183, pp.972–984.

Hayashi, T., Hou, Y., Glasser, M.F., Autio, J.A., Knoblauch, K., Inoue-Murayama, M., Coalson, T., Yacoub, E., Smith, S., Kennedy, H. and Van Essen, D.C., 2021. The nonhuman primate neuroimaging and neuroanatomy project. NeuroImage, 229, p.117726.

Ibrahim, T.S., Lee, R., Abduljalil, A.M., Baertlein, B.A. and Robitaille, P.M.L., 2001. Dielectric resonances and B1 field inhomogeneity in UHFMRI: computational analysis and experimental findings. Magnetic resonance imaging, 19(2), pp.219–226.

Jiru, F. and Klose, U., 2006. Fast 3D radiofrequency field mapping using echo-planar imaging. Magnetic Resonance in Medicine: An Official Journal of the International Society for Magnetic Resonance in Medicine, 56(6), pp.1375–1379.

Kiefer, J., 1953. Sequential minimax search for a maximum. Proceedings of the American mathematical society, 4(3), pp.502–506.

Kwon, D., Pfefferbaum, A., Sullivan, E.V. and Pohl, K.M., 2020. Regional growth trajectories of cortical myelination in adolescents and young adults: longitudinal validation and functional correlates. Brain imaging and behavior, 14(1), pp.242–266.

Lee, K., Cherel, M., Budin, F., Gilmore, J., Consing, K.Z., Rasmussen, J., Wadhwa, P.D., Entringer, S., Glasser, M.F., Van Essen, D.C. and Buss, C., 2015, March. Early postnatal myelin content estimate of white matter via T1w/T2w Ratio. In Medical Imaging 2015: Biomedical Applications in Molecular, Structural, and Functional Imaging (Vol. 9417, p. 94171R). International Society for Optics and Photonics.

Li, X., Zhu, Q., Janssens, T., Arsenault, J.T. and Vanduffel, W., 2019. In vivo identification of thick, thin, and pale stripes of macaque area V2 using submillimeter resolution (f) MRI at 3 T. Cerebral Cortex, 29(2), pp.544–560.

Liu, H., Ljungberg, E., Dvorak, A.V., Lee, L.E., Yik, J.T., MacMillan, E.L., Barlow, L., Li, D.K., Traboulsee, A., Kolind, S.H. and Kramer, J.L., 2020. Myelin water fraction and intra/extracellular water geometric mean T2 normative atlases for the cervical spinal cord from 3T MRI. Journal of Neuroimaging, 30(1), pp.50–57.

Liu, J., Xia, M., Wang, X., Liao, X. and He, Y., 2020. The spatial organization of the chronnectome associates with cortical hierarchy and transcriptional profiles in the human brain. Neuroimage, 222, p.117296.

Lutti, A., Dick, F., Sereno, M.I. and Weiskopf, N., 2014. Using high-resolution quantitative mapping of R1 as an index of cortical myelination. Neuroimage, 93, pp.176–188.

MacLennan, T., Seres, P., Rickard, J., Stolz, E., Beaulieu, C. and Wilman, A.H., 2021. Characterization of B1+ field variation in brain at 3 T using 385 healthy individuals across the lifespan. Magnetic Resonance in Medicine.

Maggioni, M.B., Krämer, M. and Reichenbach, J.R., 2021. Optimized gradient spoiling of UTE VFA-AFI sequences for robust T1 estimation with B1-field correction. Magnetic Resonance Imaging, 82, pp.1–8.

Mangeat, G., Govindarajan, S.T., Mainero, C. and Cohen-Adad, J., 2015. Multivariate combination of magnetization transfer, T2* and B0 orientation to study the myelo-architecture of the in vivo human cortex. NeuroImage, 119, pp.89–102.

Marques, J.P., Khabipova, D. and Gruetter, R., 2017. Studying cyto and myeloarchitecture of the human cortex at ultra-high field with quantitative imaging: R1, R2* and magnetic susceptibility. NeuroImage, 147, pp.152–163.

Marques, J.P. and Norris, D.G., 2018. How to choose the right MR sequence for your research question at 7 T and above?. NeuroImage, 168, pp.119–140.

Mars, R.B., Sotiropoulos, S.N., Passingham, R.E., Sallet, J., Verhagen, L., Khrapitchev, A.A., Sibson, N. and Jbabdi, S., 2018. Whole brain comparative anatomy using connectivity blueprints. Elife, 7, p.e35237.

Ma, Z. and Zhang, N., 2017. Cross-population myelination covariance of human cerebral cortex. Human brain mapping, 38(9), pp.4730–4743.

Metere, R., Kober, T., Möller, H.E. and Schäfer, A., 2017. Simultaneous quantitative MRI mapping of T 1, T 2* and magnetic susceptibility with multi-echo MP2RAGE. PloS one, 12(1), p.e0169265.

Milchenko, M. and Marcus, D., 2013. Obscuring surface anatomy in volumetric imaging data. Neuroinformatics, 11(1), pp.65–75.

Möller, H.E., Bossoni, L., Connor, J.R., Crichton, R.R., Does, M.D., Ward, R.J., Zecca, L., Zucca, F.A. and Ronen, I., 2019. Iron, myelin, and the brain: neuroimaging meets neurobiology. Trends in neurosciences, 42(6), pp.384–401.

Mugler III, J.P., 2014. Optimized three-dimensional fast-spin-echo MRI. Journal of magnetic resonance imaging, 39(4), pp.745–767.

Mugler III, J.P., Bao, S., Mulkern, R.V., Guttmann, C.R., Robertson, R.L., Jolesz, F.A. and Brookeman, J.R., 2000. Optimized single-slab three-dimensional spin-echo MR imaging of the brain. Radiology, 216(3), pp.891–899.

Mugler III, J.P. and Brookeman, J.R., 1990. Three-dimensional magnetization-prepared rapid gradient-echo imaging (3D MP RAGE). Magnetic resonance in medicine, 15(1), pp.152–157.

Nakamura, K., Chen, J.T., Ontaneda, D., Fox, R.J. and Trapp, B.D., 2017. T1-/T2-weighted ratio differs in demyelinated cortex in multiple sclerosis. Annals of neurology, 82(4), pp.635–639.

Nehrke, K., 2009. On the steady-state properties of actual flip angle imaging (AFI). Magnetic Resonance in Medicine: An Official Journal of the International Society for Magnetic Resonance in Medicine, 61(1), pp.84–92.

Nerland, S., Jorgensen, K.N., Nordhoy, W., Maximov, I.I., Bugge, R.A.B., Westlye, L.T., Andreassen, O.A., Geier, O.M., and Agartz, I., 2021. Multisite reproducibility and test-retest reliability of the T1w/T2w-ratio: A comparison of processing methods. Neuroimage.

Norbom, L.B., Rokicki, J., Alnæs, D., Kaufmann, T., Doan, N.T., Andreassen, O.A., Westlye, L.T. and Tamnes, C.K., 2020. Maturation of cortical microstructure and cognitive development in childhood and adolescence: a T1w/T2w ratio MRI study. Human brain mapping, 41(16), pp.4676–4690.

Oh, S.H., Kim, Y.B., Cho, Z.H. and Lee, J., 2013. Origin of B0 orientation dependent R2*(= 1/T2*) in white matter. Neuroimage, 73, pp.71–79.

Operto, G., Molinuevo, J.L., Cacciaglia, R., Falcon, C., Brugulat-Serrat, A., Suárez-Calvet, M., Grau-Rivera, O., Bargalló, N., Morán, S., Esteller, M. and Gispert, J.D., 2019. Interactive effect of age and APOE-ε4 allele load on white matter myelin content in cognitively normal middle-aged subjects. NeuroImage: Clinical, 24, p.101983.

Paquola, C., Seidlitz, J., Benkarim, O., Royer, J., Klimes, P., Bethlehem, R.A., Larivière, S., de Wael, R.V., Rodríguez-Cruces, R., Hall, J.A. and Frauscher, B., 2020. A multi-scale cortical wiring space links cellular architecture and functional dynamics in the human brain. PLoS biology, 18(11), p.e3000979.

Patel, Y., Shin, J., Drakesmith, M., Evans, J., Pausova, Z. and Paus, T., 2020. Virtual histology of multi-modal magnetic resonance imaging of cerebral cortex in young men. NeuroImage, 218, p.116968.

Qiu, Y., She, S., Zhang, S., Wu, F., Liang, Q., Peng, Y., Yuan, H., Ning, Y., Wu, H. and Huang, R., 2021. Cortical myelin content mediates differences in affective temperaments. Journal of Affective Disorders, 282, pp.1263–1271.

Reuter, M., Tisdall, M.D., Qureshi, A., Buckner, R.L., van der Kouwe, A.J. and Fischl, B., 2015. Head motion during MRI acquisition reduces gray matter volume and thickness estimates. Neuroimage, 107, pp.107–115.

Righart, R., Biberacher, V., Jonkman, L.E., Klaver, R., Schmidt, P., Buck, D., Berthele, A., Kirschke, J.S., Zimmer, C., Hemmer, B. and Geurts, J.J., 2017. Cortical pathology in multiple sclerosis detected by the T 1/T 2-weighted ratio from routine magnetic resonance imaging. Annals of neurology, 82(4), pp.519–529.

Robinson, E.C., Garcia, K., Glasser, M.F., Chen, Z., Coalson, T.S., Makropoulos, A., Bozek, J., Wright, R., Schuh, A., Webster, M. Hutter, J., Price, A., Grande, L.C., Hughes, E., Tusor, N., Bayly, P.V., Van Essen, D.C., Smith, S.M., Edwards, A.D., Hajnal, J., Jenkinson, M., Glocker, B., and Rueckert, D., 2018. Multimodal surface matching with higher-order smoothness constraints. Neuroimage, 167, pp.453–465.

Robinson, E.C., Jbabdi, S., Glasser, M.F., Andersson, J., Burgess, G.C., Harms, M.P., Smith, S.M., Van Essen, D.C. and Jenkinson, M., 2014. MSM: a new flexible framework for Multimodal Surface Matching. Neuroimage, 100, pp.414–426.

Rowley, C.D., Tabrizi, S.J., Scahill, R.I., Leavitt, B.R., Roos, R.A., Durr, A. and Bock, N.A., 2018. Altered Intracortical T1-weighted/T2-weighted ratio signal in Huntington’s disease. Frontiers in neuroscience, 12, p.805.

Sacolick, L.I., Wiesinger, F., Hancu, I. and Vogel, M.W., 2010. B1 mapping by Bloch-Siegert shift. Magnetic resonance in medicine, 63(5), pp.1315–1322.

Salimi-Khorshidi, G., Douaud, G., Beckmann, C.F., Glasser, M.F., Griffanti, L. and Smith, S.M., 2014. Automatic denoising of functional MRI data: combining independent component analysis and hierarchical fusion of classifiers. Neuroimage, 90, pp.449–468.

Sánchez-Panchuelo, R.M., Besle, J., Mougin, O., Gowland, P., Bowtell, R., Schluppeck, D. and Francis, S., 2014. Regional structural differences across functionally parcellated Brodmann areas of human primary somatosensory cortex. Neuroimage, 93, pp.221–230.

Seifert, A.C., Umphlett, M., Hefti, M., Fowkes, M. and Xu, J., 2019. Formalin tissue fixation biases myelin-sensitive MRI. Magnetic resonance in medicine, 82(4), pp.1504–1517.

Sereno, M.I., Lutti, A., Weiskopf, N. and Dick, F., 2013. Mapping the human cortical surface by combining quantitative T 1 with retinotopy. Cerebral cortex, 23(9), pp.2261–2268.

Shafee, R., Buckner, R.L. and Fischl, B., 2015. Gray matter myelination of 1555 human brains using partial volume corrected MRI images. Neuroimage, 105, pp.473–485.

Shams, Z., Norris, D.G. and Marques, J.P., 2019. A comparison of in vivo MRI based cortical myelin mapping using T1w/T2w and R1 mapping at 3T. PloS one, 14(7), p.e0218089.

Shatil, A.S., Uddin, M.N., Matsuda, K.M. and Figley, C.R., 2018. Quantitative ex vivo MRI changes due to progressive formalin fixation in whole human brain specimens: longitudinal characterization of diffusion, relaxometry, and myelin water fraction measurements at 3T. Frontiers in medicine, 5, p.31.

Sled, J.G. and Pike, G.B., 1998. Standing-wave and RF penetration artifacts caused by elliptic geometry: an electrodynamic analysis of MRI. IEEE transactions on medical imaging, 17(4), pp.653–662.

Smith, S.M., Beckmann, C.F., Andersson, J., Auerbach, E.J., Bijsterbosch, J., Douaud, G., Duff, E., Feinberg, D.A., Griffanti, L., Harms, M.P., Kelly, M., Laumann, T., Miller, K.L., Moeller, S., Petersen, S., Power, J., Salimi-Khorshidi, G., Snyder, A.Z., Vu, A.T., Woolrich, M.W., Xu, J., Yacoub, E., Ugurbil, K., Van Essen, D.C. and Glasser, M.F. 2013. Resting-state fMRI in the human connectome project. Neuroimage, 80, pp.144–168.

Smith, S.M., Jenkinson, M., Johansen-Berg, H., Rueckert, D., Nichols, T.E., Mackay, C.E., Watkins, K.E., Ciccarelli, O., Cader, M.Z., Matthews, P.M. and Behrens, T.E., 2006. Tract-based spatial statistics: voxelwise analysis of multi-subject diffusion data. Neuroimage, 31(4), pp.1487–1505.

Somerville, L.H., Bookheimer, S.Y., Buckner, R.L., Burgess, G.C., Curtiss, S.W., Dapretto, M., Elam, J.S., Gaffrey, M.S., Harms, M.P., Hodge, C. and Kandala, S., 2018. The Lifespan Human Connectome Project in Development: A large-scale study of brain connectivity development in 5-21 year olds. Neuroimage, 183, pp.456–468.

Stikov, N., Campbell, J.S., Stroh, T., Lavelée, M., Frey, S., Novek, J., Nuara, S., Ho, M.K., Bedell, B.J., Dougherty, R.F. and Leppert, I.R., 2015. In vivo histology of the myelin g-ratio with magnetic resonance imaging. Neuroimage, 118, pp.397–405.

Stüber, C., Morawski, M., Schäfer, A., Labadie, C., Wähnert, M., Leuze, C., Streicher, M., Barapatre, N., Reimann, K., Geyer, S. and Spemann, D., 2014. Myelin and iron concentration in the human brain: a quantitative study of MRI contrast. Neuroimage, 93, pp.95–106.

Sun, H., Cleary, J.O., Glarin, R., Kolbe, S.C., Ordidge, R.J., Moffat, B.A. and Pike, G.B., 2020. Extracting more for less: multi-echo MP2RAGE for simultaneous T1-weighted imaging, T1 mapping, mapping, SWI, and QSM from a single acquisition. Magnetic resonance in medicine, 83(4), pp.1178–1191.

Teraguchi, M., Yamada, H., Yoshida, M., Nakayama, Y., Kondo, T., Ito, H., Terada, M. and Kaneoke, Y., 2014. Contrast enrichment of spinal cord MR imaging using a ratio of T1-weighted and T2-weighted signals. Journal of Magnetic Resonance Imaging, 40(5), pp.1199–1207.

Thompson, E., Mohammadi-Nejad, A.R., Robinson, E.C., Andersson, J.L., Jbabdi, S., Glasser, M.F., Bastiani, M. and Sotiropoulos, S.N., 2020. Non-negative data-driven mapping of structural connections with application to the neonatal brain. NeuroImage, 222, p.117273.

Tisdall, M.D., Hess, A.T., Reuter, M., Meintjes, E.M., Fischl, B. and van der Kouwe, A.J., 2012. Volumetric navigators for prospective motion correction and selective reacquisition in neuroanatomical MRI. Magnetic resonance in medicine, 68(2), pp.389–399.

Toschi, N. and Passamonti, L., 2019. Intra-cortical myelin mediates personality differences. Journal of personality, 87(4), pp.889–902.

Trampel, R., Bazin, P.L., Pine, K. and Weiskopf, N., 2019. In-vivo magnetic resonance imaging (MRI) of laminae in the human cortex. Neuroimage, 197, pp.707–715.

Turner, R., 2019. Myelin and modeling: Bootstrapping cortical microcircuits. Frontiers in neural circuits, 13, p.34.

Uddin, M.N., Figley, T.D., Marrie, R.A., Figley, C.R. and CCOMS Study Group, 2018. Can T1w/T2w ratio be used as a myelin-specific measure in subcortical structures? Comparisons between FSE-based T1w/T2w ratios, GRASE-based T1w/T2w ratios and multi-echo GRASE-based myelin water fractions. NMR in Biomedicine, 31(3), p.e3868.

Uddin, M.N., Figley, T.D., Solar, K.G., Shatil, A.S. and Figley, C.R., 2019. Comparisons between multi-component myelin water fraction, T1w/T2w ratio, and diffusion tensor imaging measures in healthy human brain structures. Scientific Reports, 9(1), pp.1–17.

Uğurbil, K., Xu, J., Auerbach, E.J., Moeller, S., Vu, A.T., Duarte-Carvajalino, J.M., Lenglet, C., Wu, X., Schmitter, S., Van de Moortele, P.F. and Strupp, J., 2013. Pushing spatial and temporal resolution for functional and diffusion MRI in the Human Connectome Project. Neuroimage, 80, pp.80–104.

Vaidya, M.V., Collins, C.M., Sodickson, D.K., Brown, R., Wiggins, G.C. and Lattanzi, R., 2016. Dependence of and field patterns of surface coils on the electrical properties of the sample and the MR operating frequency. Concepts in Magnetic Resonance Part B: Magnetic Resonance Engineering, 46(1), pp.25–40.

Van de Moortele, P.F., Auerbach, E.J., Olman, C., Yacoub, E., Uğurbil, K. and Moeller, S., 2009. T1 weighted brain images at 7 Tesla unbiased for Proton Density, T2* contrast and RF coil receive B1 sensitivity with simultaneous vessel visualization. Neuroimage, 46(2), pp.432–446.

van der Kouwe, A.J., Benner, T., Salat, D.H. and Fischl, B., 2008. Brain morphometry with multiecho MPRAGE. Neuroimage, 40(2), pp.559–569.

Van Essen, D.C., Glasser, M.F., Dierker, D.L., Harwell, J. and Coalson, T., 2012. Parcellations and hemispheric asymmetries of human cerebral cortex analyzed on surface-based atlases. Cerebral cortex, 22(10), pp.2241–2262.

Van Essen, D.C., Smith, J., Glasser, M.F., Elam, J., Donahue, C.J., Dierker, D.L., Reid, E.K., Coalson, T. and Harwell, J., 2017. The brain analysis library of spatial maps and atlases (BALSA) database. Neuroimage, 144, pp.270–274.

Van Essen, D.C., Smith, S.M., Barch, D.M., Behrens, T.E., Yacoub, E., Ugurbil, K. and Wu-Minn HCP Consortium, 2013. The WU-Minn human connectome project: an overview. Neuroimage, 80, pp.62–79.

Vidal-Piñeiro, D., Walhovd, K.B., Storsve, A.B., Grydeland, H., Rohani, D.A. and Fjell, A.M., 2016. Accelerated longitudinal gray/white matter contrast decline in aging in lightly myelinated cortical regions. Human brain mapping, 37(10), pp.3669–3684.

Von Bonin G (1950) Essay on the cerebral cortex. Springfield, IL: Charles C Thomas.

Wang, F., Dong, Z., Reese, T.G., Rosen, B., Wald, L.L. and Setsompop, K., 2022. 3D Echo Planar Time-resolved Imaging (3D-EPTI) for ultrafast multi-parametric quantitative MRI. NeuroImage, 250, p.118963.

Wang, D., Heberlein, K., LaConte, S. and Hu, X., 2004. Inherent insensitivity to RF inhomogeneity in FLASH imaging. Magnetic Resonance in Medicine: An Official Journal of the International Society for Magnetic Resonance in Medicine, 52(4), pp.927–931.

Wang, J., Qiu, M. and Constable, R.T., 2005. In vivo method for correcting transmit/receive nonuniformities with phased array coils. Magnetic Resonance in Medicine: An Official Journal of the International Society for Magnetic Resonance in Medicine, 53(3), pp.666–674.

Wei, W., Zhang, Y., Li, Y., Meng, Y., Li, M., Wang, Q., Deng, W., Ma, X., Palaniyappan, L., Zhang, N. and Li, T., 2020. Depth-dependent abnormal cortical myelination in first-episode treatment-naïve schizophrenia. Human brain mapping, 41(10), pp.2782–2793.

Weiskopf, N., Lutti, A., Helms, G., Novak, M., Ashburner, J. and Hutton, C., 2011. Unified segmentation based correction of R1 brain maps for RF transmit field inhomogeneities (UNICORT). Neuroimage, 54(3), pp.2116–2124.

Weiskopf, N., Suckling, J., Williams, G., Correia, M.M., Inkster, B., Tait, R., Ooi, C., Bullmore, E.T. and Lutti, A., 2013. Quantitative multi-parameter mapping of R1, PD*, MT, and R2* at 3T: a multi-center validation. Frontiers in neuroscience, 7, p.95.

Yang, G., Bozek, J., Han, M. and Gao, J.H., 2020. Constructing and evaluating a cortical surface atlas and analyzing cortical sex differences in young Chinese adults. Human brain mapping, 41(9), pp.2495–2513.

Yarnykh, V.L., 2007. Actual flip-angle imaging in the pulsed steady state: a method for rapid three-dimensional mapping of the transmitted radiofrequency field. Magnetic Resonance in Medicine: An Official Journal of the International Society for Magnetic Resonance in Medicine, 57(1), pp.192–200.

Yarnykh, V.L., 2016. Time-efficient, high-resolution, whole brain three-dimensional macromolecular proton fraction mapping. Magnetic resonance in medicine, 75(5), pp.2100–2106.

Zaretskaya, N., Fischl, B., Reuter, M., Renvall, V. and Polimeni, J.R., 2018. Advantages of cortical surface reconstruction using submillimeter 7 T MEMPRAGE. Neuroimage, 165, pp.11–26.

Zhang, H., Schneider, T., Wheeler-Kingshott, C.A. and Alexander, D.C., 2012. NODDI: practical in vivo neurite orientation dispersion and density imaging of the human brain. Neuroimage, 61(4), pp.1000–1016.

